# Heparin promotes fibrillation of most phenol soluble modulin peptides from *S. aureus*: a possible strengthening of the bacterial biofilm

**DOI:** 10.1101/2021.03.07.434294

**Authors:** Zahra Najarzadeh, Masihuz Zaman, Vita Serekaité, Kristian Strømgaard, Maria Andreasen, Daniel E. Otzen

## Abstract

Phenol soluble modulins (PSMs) are virulence peptides secreted by different *Staphylococcus aureus* strains. In addition, PSMs are able to form amyloid fibrils which may strengthen the biofilm matrix. The highly sulfated glycosaminoglycan heparin promotes *S.aureus* infection but the basis for this is unclear. We hypothesized that heparin promotes PSM fibrillation and in this way aids bacterial colonization. Here we address this hypothesis using a combination of different biophysical techniques along with peptide microarrays. We find that heparin accelerates fibrillation of all α-PSMs (except PSMα2) and δ-toxin, but inhibits β-PSMs’ fibrillation by blocking nucleation. Given that *S. aureus* secretes higher levels of α-PSMs than β-PSMs peptidess, heparin is likely to overall promote fibrillation. Heparin binding is driven by multiple positively charged lysine residues in α-PSMs and δ-toxins, whose removal strongly reduces affinity. Binding of heparin does not alter the final fibril conformation. Rather, heparin provides a scaffold to catalyze or inhibit fibrillation. Our findings suggest that heparin may strengthen bacterial biofilm through increased PSM fibrillation.

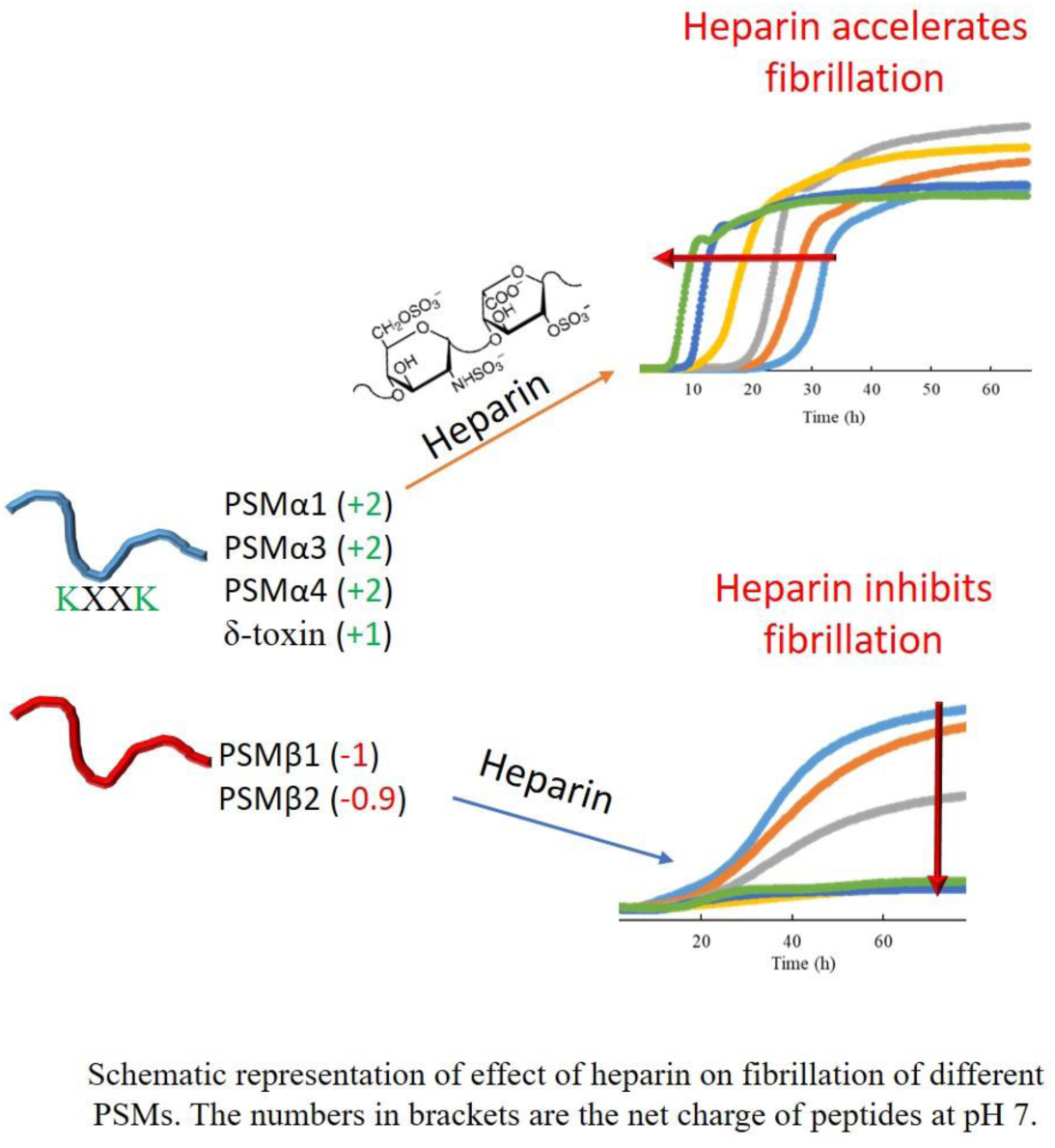

## INTRODUCTION

Functional bacterial amyloids are proteins secreted from bacteria which self-assemble to form highly ordered β-sheet-rich fibrils or amyloids. These fibrils promote bacterial biofilm formation by acting as structural scaffolds in the biofilm matrix, leading to increased antibiotic resistance [1]. The most well-understood functional bacterial amyloids are curli in *Escherichia coli* [2], Fap in *Pseudomonas* [3], TasA in *Bacillus subtilis* [4] and phenol-soluble modulins (PSMs) in *Staphylococcus* strains [5]. Unlike the other amyloids, PSMs are short peptides with multiple functions. Besides their ability to strengthen biofilm through amyloid formation, they act as virulence factors which lyse neutrophils and erythrocytes and stimulate inflammatory responses [6–8]. As amphipathic peptides they are both surface- and membrane-active and this is thought to lead to cell permeabilization, as well as encouraging early steps in biofilm formation [5, 9]. PSMs are classified according to their length: the shortest (20-25 residues) and most abundant are the four α-PSMs and δ-toxin which all adopt an α-helical amphipathic structure in solution. The two β-PSMs (∼44 residues), both containing a C-terminal amphipathic α-helix, are found in much lower amounts *in vivo* [10].

Extracellular fibrils isolated from *S. aureus* biofilm contain several different PSMs [5]. Almost all individual PSMs fibrillate in the classic cross-β amyloid motif with β-strands perpendicular to the fibril axis. The only exceptions are PSMα2 and δ-toxin, which do not fibrillate on their own, and PSMα3, which is the first reported example of the cross-α fold, in which monomeric α-helices are oriented perpendicularly to the fibril axis [9, 11]. Preformed PSMs fibrils (particulary those formed by PSMα1) accelerate fibrillation of other PSM peptides and even seed fibrillation of PSMα2 and δ-toxin [11]. PSMα3 fibrillates very rapidly and thus provides seeds to promote fibrillation of PSMs such as PSMα1 (despite the difference in structure) which in turn is very efficient at accelerating the fibrillation of other PSMs [11].

Besides inter-PSM interactions, other components found in the biofilm matrix such as polysaccharides, proteins and extracellular DNA (eDNA) may influence fibrillation. eDNA is known to promote PSMα1 fibrillation [12]; similarly, bacterial biosurfactants such as rhamnolipids and outer-membrane lipopolysaccharides generally promote amyloid formation [13]. Importantly, eukaryotic host factors can also play a role. Chief among these is heparin, a glycosaminoglycan which thanks to its many sulfate and carboxyl groups is the most highly anionic biomacromolecule known [14]. Heparin is normally stored intracellularly in secretory granules and released upon tissue injury to act as an anticoagulant, preventing clot formation by fibrinogen [15, 16]. This has inspired its use as an anticoagulant in patient catheters, especially for kidney dialysis. In a bacterial context, however, heparin stimulates *S. aureus* biofilm formation by accumulating in the biofilm matrix, probably by binding to cell-surface proteins as a mimic of extracellular DNA. This often leads to catheter infections [17–19]. *In vitro* heparin effectively induces fibrillation of a range of amyloidogenic proteins e.g lysozyme, α-synuclein, transthyretin, β_2_-microglobulin, Aβ and the prion protein [20–24]. These observations prompted us to hypothesize that heparin might encourage biofilm formation through PSM fibrillation. Here we investigate how heparin affects PSM fibrillation processes by a combination of biophysical techniques, peptide arrays and biofilm formation assays.

## MATERIALS AND METHODS

### Peptides, reagents and solutions

The peptides PSMα1 (MGIIAGIIKVIKSLIEQFTGK), PSMα2 (MGIIAGIIKFIKGLIEKFTGK), PSMα3 (MEFVAKLFKFFKDLLGKFLGNN), PSMα4 (MAIVGTIIKIIKAIIDIFAK), PSMβ1 (MEGLFNAIKD TVTAAINNDG AKLGTSIVSI VENGVGLLGK LFGF), PSMβ2 (MTGLAEAIAN TVQAAQQHDS VKLGTSIVDI VANGVGLLGK LFGF) and δ-toxin (MAQDIISTIG DLVKWIIDTV NKFTKK) were purchased from GenScript Biotech, Netherlands. All peptides were N-terminally formylated with a purity of >95 %. All reagents and chemicals were of analytical grade. Heparin-fluorescein conjugate, and chemicals including Hexafluoroisopropanol (HFIP), Thioflavin T, trifluoroacetic acid (TFA) and crystal violet solution (2.3%) were from Sigma-Aldrich Ltd. Heparin (Cat # Y0001282) was from European Pharmacopoeia, Europe. Dimethyl-sulfoxide (DMSO) was from Merck. Peptide stock solutions were filtered using PVDF 0.22 µm syringe filters (Millipore Milex-HV) before use.

### Peptide pre-treatment

For aggregation kinetics and secondary structure analysis (CD and FTIR), each PSM peptide stock was pretreated to disassemble any pre-formed aggregates. All seven dry lyophilized peptides (PSMα1-4, PSMβ1-2 and δ-toxin) were freshly dissolved to a final concentration of 0.5 mg/mL in HFIP:TFA (1:1 v/v) and sonicated for 5×x20 seconds with 30 seconds intervals using a probe sonicator, followed by incubation at room temperature for 1 h. Solutions were then aliquoted out and organic solvent was evaporated using a speedvac (1000 rpm for 3-4 h) at room temperature. Dried peptide stocks were stored at -80 °C prior to use.

### Thioflavin T (ThT) Fibrillation Assay

10 mg of heparin was dissolved in 1 mL milliQ water and passed through a PVDF 0.45 µm syringe filter. PSMs were thawed and dissolved in dimethyl sulfoxide (DMSO) to 10 mg/mL prior to use. All seven freshly prepared peptides were diluted (typically to 0.25-1 mg/mL) into sterile milliQ water containing 40 µM ThT with 0-1 mg/mL heparin in a final volume of 100 μl in a 96 well black polystyrene microtiter plates. Different PSMs with a fixed monomeric peptide concentration (0.25 mg/mL to 1.0 mg/mL) were supplemented with appropriate amounts of heparin (0-1 mg/mL) for the entire study. The low residual amounts of DMSO and high amount of heparin itself from the stock solution did not give rise to any fibrillation (Fig. S1F). ThT fluorescence was monitored on a Fluostar Omega (BMG Labtech, Germany) plate reader in bottom reading mode at 37 °C under quiescent conditions. The plate was sealed with metal sealing tape to prevent evaporation. ThT fluorescence of all PSMs except PSMα3 was measured every 10 minutes with an excitation filter of 450 nm and an emission filter of 482 nm under quiescent conditions. For PSMα3, ThT fluorescence was measured every 20 s with an excitation filter of 450 nm and an emission filter of 482 nm. 0-1 mg/mL heparin alone was tested in separate experiments. All measurements were in triplicate.

### Synchrotron radiation circular dichroism spectroscopy (SRCD)

SRCD spectra of PSM fibrils were collected at the AU-CD beamline of the ASTRID2 synchrotron, Aarhus University, Denmark. PSMs samples aggregated in the absence and presence of heparin were collected directly from the 96-well plates and pelleted at 13 krpm for 30 min. The supernatant was gently removed from each sample, and the pellet fraction was re-suspended in milliQ water. Three to five successive spectra of fibrillated PSMs (in the absence and presence of heparin) were recorded from 280 to 170 nm in a 0.2 mm path length cuvette with a dwell time of 2 sec at 1 nm intervals at 25 °C. All SRCD spectra were processed and their respective averaged baseline (a solution containing all components of the sample, except the protein) subtracted, smoothing with a 7 pt Savitzky-Golay filter. The secondary structural content of individual SRCD spectra of PSM fibrils samples (in the absence and presence of heparin) was determined using DichroWeb [25, 26]. Each spectrum was fitted using three different analysis programs (Selecon3, Contin and CDSSTR) with the SP175 reference data set [27]. An average of the structural component contributions from the three analysis programs was used.

### Thermal fibril stability by Circular dichroism analysis

Thermal CD spectra were recorded on a JASCO-810 (Jasco Spectroscopic Co., Ltd., Hachioji City, Japan) spectrophotometer equipped with a Peltier thermally controlled cuvette holder. At the end of ThT kinetics experiments, individual triplicate samples fibrillated in the absence and presence of the maximum concentrations of heparin i.e., 3 µg/mL for PSMα1, 40 µg/mL for PSMα3, 50 µg/mL for PSMα4, 250 µg/mL for PSMβ1, 1 mg/mL for PSMβ2 and 1 mg/mL for δ-toxin were pelleted at 13 kpm for 30 min and supernatant was removed. The remaining pellets were re-suspended in the same volume of milliQ water and thermal scans were recorded at 220 nm from 25 to 95 °C with a step size of 0.1 °C.

### Attenuated Total Reflectance Fourier Transform Infrared Spectroscopy (ATR-FTIR)

PSM fibrils were prepared as above. 5 µL of sample was applied on the surface of the ATR module and dried under a steam of nitrogen gas. FTIR spectra were recorded on a Tensor 27 FTIR instrument (Bruker optics, Billerica, Massachusetts, USA) equipped with an attenuated total reflection accessory with a continuous flow of N_2_ gas. Measurements were performed as an accumulation of 64 scans with a spectral resolution of 2 cm^-1^ over a range of 1000 to 3998 cm^-1^. Atmospheric compensation and baseline correlation was executed. The individual components of the spectrum were determined through second derivative analysis of curves by employing OPUS 5.5 software (Bruker, Billerica, Massachusetts, USA). For comparative studies, all absorbance spectra were normalized.

### Transmission electron microscopy (TEM)

The morphology of the PSMs species in the presence and absence of heparin was analyzed with TEM. 5 μL of endpoint samples from the non-shaking ThT assay of each PSM was transferred to carbon-coated formvar electron microscopy grids followed by two minute incubation at room temperature. Buffer was then removed by blotting the grid with Whatman filter paper, washed with 5 µL milliQ water and stained with 1% uranyl acetate for 2 min, after which excess staining solution was removed with filter paper. Finally, the grids were washed twice with 5 μL of milliQ and dried before analysis. Samples were viewed in a Morgagni 268 FEI Phillips Electron microscope equipped with a CCD digital camera, operated at 80 kV.

### Heparin-PSM interactions measured using a peptide array

To probe interactions between heparin and PSMs, 351 different 10-residue peptides corresponding to different parts of the PSM sequences were immobilized on a microarray chip and incubated with fluorescein-labeled heparin [28]. In this array (full list provided in Table S3 and S4), each new peptide constituted a 10-residue window of a given PSM sequence shifted forward by 1 residue compared to the preceding peptide, giving a 9-residue overlap. For Ala scanning, each residue in a given 10-residue sequence was consecutively replaced by Ala before moving on to the next 10-residue peptide. As part of this procedure, the microarray was first blocked in a solution containing 3 % (w/v) whey protein in Tris saline buffer with 0.1 % Tween-20 (TSB-T) incubated overnight at 4 °C and washed three times with TSB-T. Subsequently it was incubated with fluorescein-labeled heparin (diluted to 0.05 mg/mL in PBS) for 4 h at room temperature. The microarray was washed 3 times with TSB-T, air-dried in the dark and scanned using a Typhoon Trio scanner (GE Life Sciences, Pittsburgh, PA). Dot intensities in the scanned image were quantified using ImageJ.

### Biofilm measurements with Crystal violet

*Staphylococcus aureus* (strain Newman) and three deletion mutants ΔPSMα, ΔPSMβ and ΔPSMα/β were grown on an LB plate overnight. A single colony was transferred to TSB medium, grown up overnight at 37 °C and then adjusted to OD ∼0.5 and further diluted 1:100 in fresh PNG media (3.3 g/L peptone; 2.6 g/L NaCl; 3.3 g/L glucose). Heparin was added (from a 5 mg/mL stock in milliQ water) to the desired concentration and 100 µL was then incubated in a 96 well plate for 24 h at 37 °C with mild shaking (50 rpm). The solutions were gently removed from all wells, after which the biofilm was washed once with PBS and air dried for 30 min at room temperature (RT). 100 µL of crystal violet solution 2.3% (Sigma) was added to all wells, incubated at RT for 10 min and removed. All wells were washed twice with PBS. The plate was air dried for 30 min, after which 150 µL of 33 % (v/v) acetic acid was added to each well to release the dye attached to the biofilm. Finally, the absorbance of the released crystal violet was measured at 590 nm.

## Results

### Aggregation kinetics of different PSMs is influenced by heparin

To investigate the effect of heparin on the fibrillation of all seven PSM peptides (PSMα1-4, PSMβ1-2 and δ-toxin), we incubated all PSMs individually (at fixed monomeric concentration) with different concentrations of heparin under quiescent conditions. Aggregation kinetics were monitored using the amyloid-binding dye ThT [29]. We have previously reported that PSMα1, PSMα3, PSMβ1 and PSMβ2 reproducibly aggregate to ThT-binding amyloid fibrils on the hour scale under quiescent conditions with different nucleating mechanism [11]. The addition of high molecular weight heparin dramatically changes the aggregation kinetics of all PSMs, but the effect varies between PSMs.

For PSMα1, as little as 1.0 μg/mL heparin increases end-level ThT fluorescence intensity and decreases the timescale for the completion of aggregation kinetics from ∼40 h to ∼25 h (Fig. 1A). Heparin also reduces the lag time from 22 h (heparin-free) to 7 hr (3.0 μg/mL heparin) in a dose-dependent manner (Fig. 1A). The lag phase decrease to 1-3 h up to 20 μg/mL heparin, above which it is completely abolished, accompanied by a reduction in ThT end point fluorescence (Fig. S1A). PSMα2 was not observed to undergo fibrillation (measured as an increase in ThT fluorescence) in the absence or presence of up to 1 mg/mL of heparin (Fig. S1B). In contrast, PSMα3 fibrillated readily and with a sigmoidal time curve both in the absence and presence of 0-50 µg/mL heparin under quiescent conditions. Heparin significantly reduced lag times and correspondingly increased end-point ThT fluorescence. Thus 40 µg/mL heparin induced a 5-fold increase in fluorescence and a ∼ 4 fold reduced lag time (Fig. 1B).

**Figure 1:**
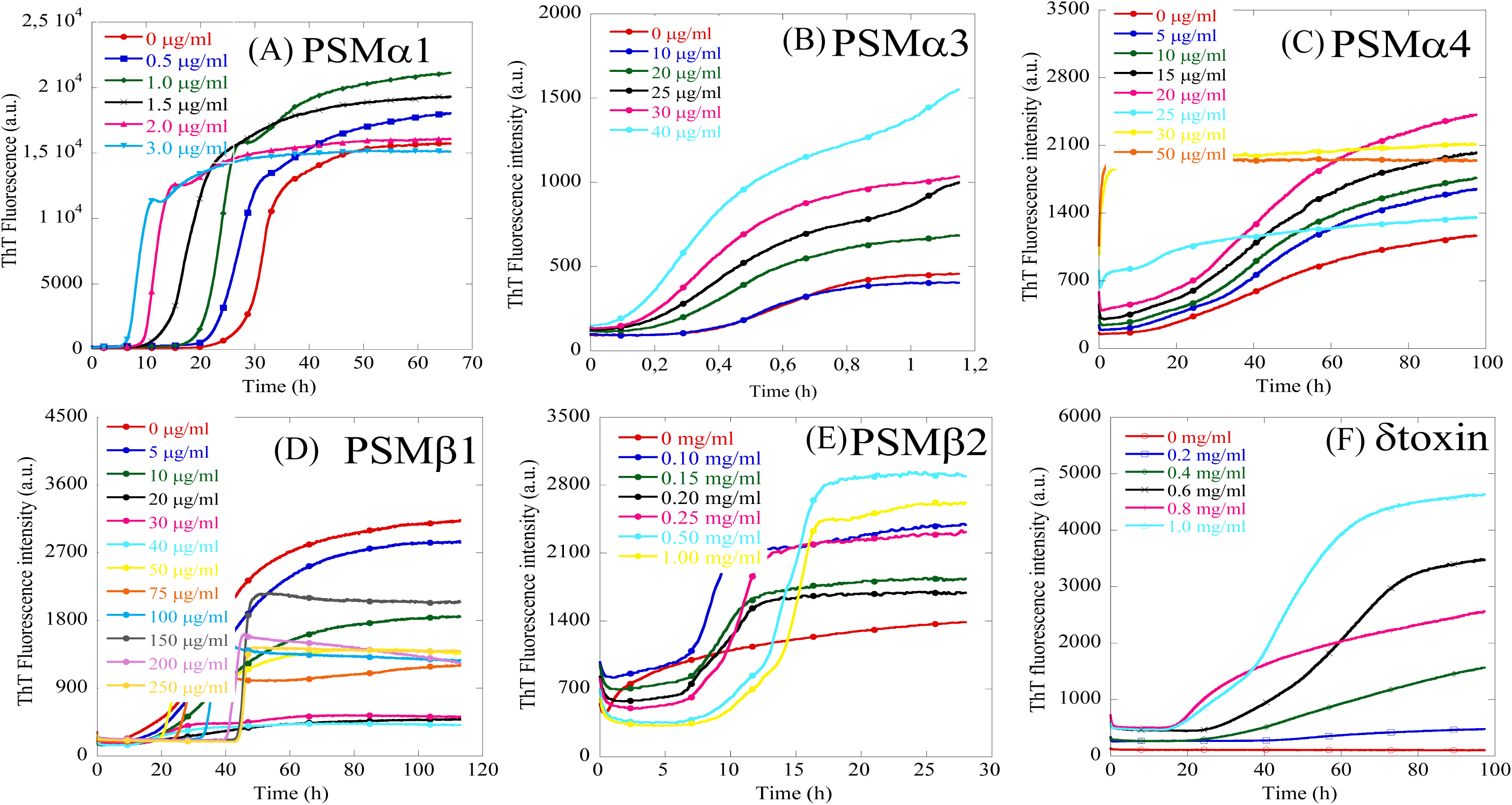
Thioflavin T time curves for PSM aggregation under quiescent conditions in the absence and presence of different heparin concentrations. The data depict representatives of triplicate experiments. PSMa1-4 and PSMb2 were incubated at 0.25 mg/mL PSM with the indicated range of heparin concentrations. PSMβ1 and δ-toxin were incubated at 0.025 and 0.3 mg/mL peptide, respectively.

The last α-PSM construct (PSMα4) shows the same ThT kinetics up to 20 μg/mL heparin though with an increase in overall ThT fluorescence (Fig. 1C). However when the ThT signals are normalized, the signals collapse to the same time curve (Fig. S1C). Above 25 µg/mL heparin, the lag time is abolished (Fig. S1D) and the data for 30-500 µg/ml heparin could be fitted with an exponential decay. The resultant rate constant (k) and amplitude of the reaction (A) decreased with [heparin] (Fig. S1E).

For PSMβ1, a more complex scenario is revealed. In the absence of heparin, the peptide fibrillates with a lag time of ∼8 h (Fig. 1D). This increases to ∼18 h in 10-40 µg/mL heparin with a major decrease in ThT fluorescence intensity (Fig. 1D). There is then an abrupt shift around 50 µg/mL where the lag time remains ∼20 h but with a much shorter and steeper elongation phase, leading to a medium level of ThT fluorescence. Increasing [heparin] above 250 µg/mL only slightly increases lag-times. PSMβ2 fibrillates with a lag time of ∼1 h in the absence of heparin. Heparin increases both the lag time and the endpoint ThT fluorescence (Fig. 1E). While δ-toxin on its own did not show any ThT fluorescence increase, heparin dramatically increased its ThT intensity with lag times decreasing from ∼45 h (0.2-0.4 mg/mL heparin) to ∼18-22 h (1 mg/mL heparin) (Fig. 1F).

To establish how heparin affects the microscopic steps during the aggregation of PSMs, we turned to the programme Amylofit [30]. Kinetic parameters from our previous analysis of PSMα1 aggregation in the absence of heparin [11] were used as fixed global parameters, while only one compound rate constant was individually fitted to each heparin concentration. This approach has previously been used for other amylogenic proteins to establish how *e.g.* inhibitors act on specific microscopic steps during aggregation [31–33]. We now go through the fits for the individual PSMs. Fits to kinetic data are shown in Fig. S2 and results from these fits in Fig. 2.

**Figure 2:**
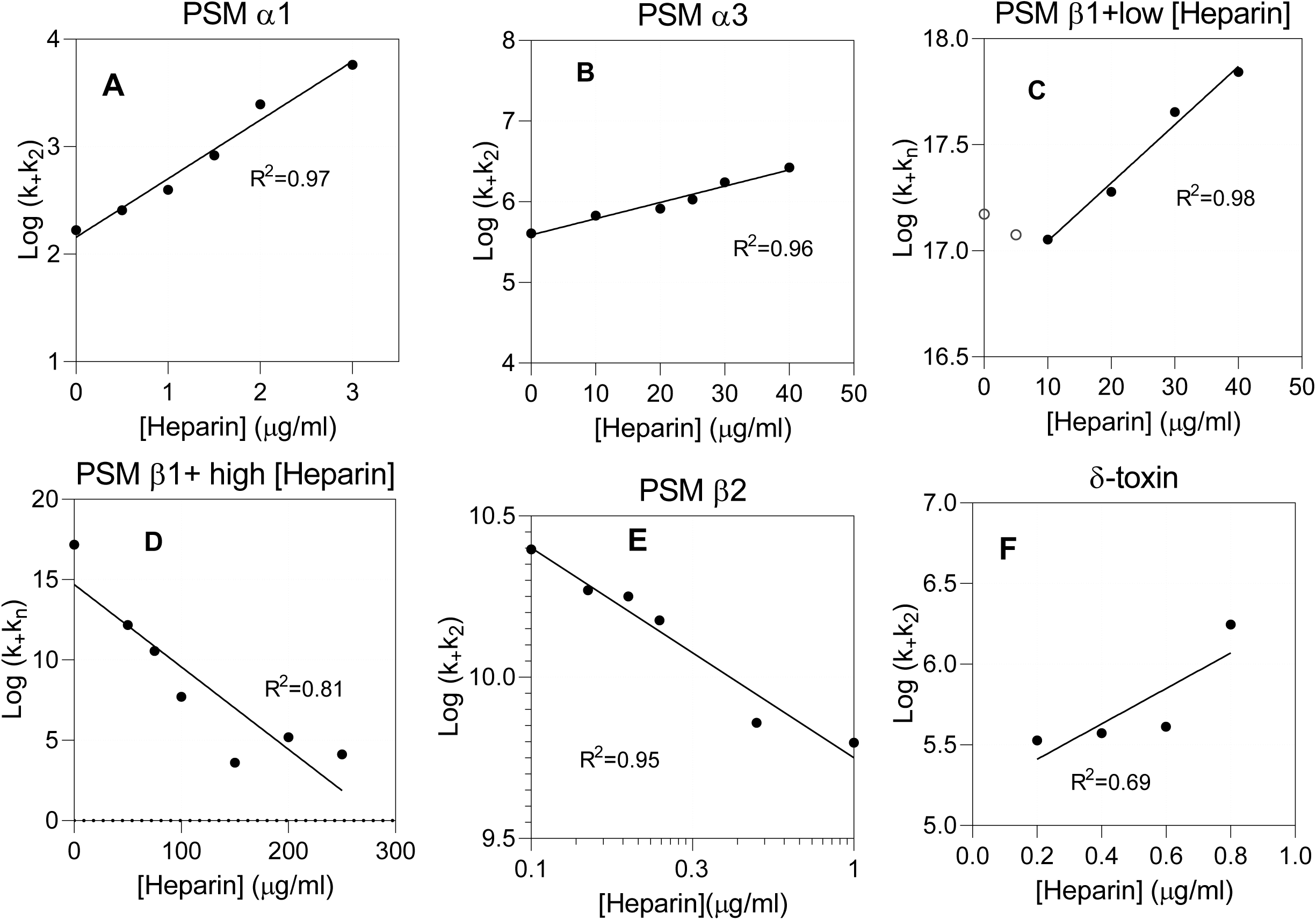
Plots of different composite rate constants (obtained from fits to PSM aggregation data) versus heparin concentration. (A) PSMα1, (B) PSMα3, (C) PSMβ1 at low heparin concentrations, (D) PSMβ1 at high heparin concentrations, (E) PSMβ2 (log-log plot), (F) δ-toxin (showing poor linear correlation with heparin concentration).

For **PSMα1**, we found the best fit when we allowed k_+_k_2_ to vary and restricted k_+_k_n_, *n*_c_ and *n*_2_ to the heparin-free values (Fig. S2A). Since k_+_k_n_ was kept constant during the data fitting, the kinetic parameter within the parameter k_+_k_2_ mostly affected by the presence of heparin is expected to be k_2_. Interestingly, log k_+_k_2_ values increase in a linear fashion when plotted against [heparin] (Fig. 2A). Similar linear relationships are seen in plots of the log of unfolding rate constants versus denaturant (urea, GdmCl) [34] or surfactant (SDS) concentrations [35]. While denaturants are present at molar concentrations (> 100 mg/mL) and rely on weak interactions with the protein, SDS effects are seen at low mM (∼1 mg/mL) concentrations and are ascribed to high affinities (driven by electrostatics) and clustering on the protein (driven by hydrophobic effects). Given the low concentrations of heparin used (∼1 µg/mL), these interactions are clearly strong and may also be cooperative in nature.

Following this pattern, the best Amylofit fits of **PSMα3** aggregation in heparin were also obtained by varying k_+_k_2_, indicating that k_2_ is most affected as is the case for PSMα1 (Fig. S2B, Table 1). This leads to a similar semi-log relationship (Fig. 2B), although the slope is reduced by a factor ∼30 (slope of semi-log plots: ∼0.54 for PSMα1 and 0.02 for PSMα3), indicating a somewhat weaker heparin-PSMα3 interaction.

**Table 1.**
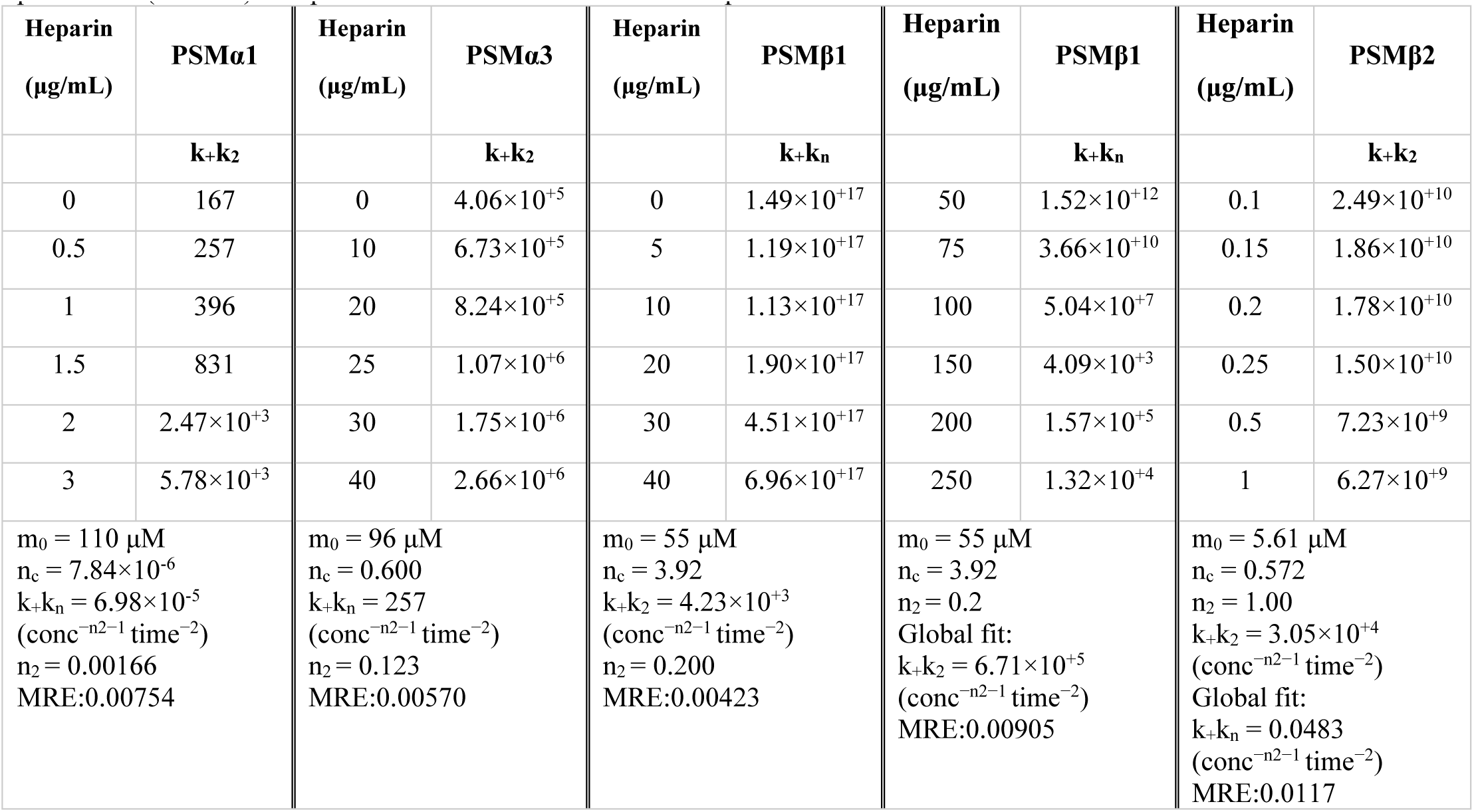
Kinetic parameters of data fitting using the webserver Amylofit in the presence of heparin using global constants for all parameters (last row) except one which is fitted to individual heparin concentrations

| Heparin<br>($\mu\text{g/mL}$) | PSMα1 | Heparin<br>($\mu\text{g/mL}$) | PSMα3 | Heparin<br>($\mu\text{g/mL}$) | PSMβ1 | Heparin<br>($\mu\text{g/mL}$) | PSMβ1 | Heparin<br>($\mu\text{g/mL}$) | PSMβ2 |
| --- | --- | --- | --- | --- | --- | --- | --- | --- | --- |
| | $k+k_2$ | | $k+k_2$ | | $k+k_n$ | | $k+k_n$ | | $k+k_2$ |
| 0 | 167 | 0 | $4.06 \times 10^{+5}$ | 0 | $1.49 \times 10^{+17}$ | 50 | $1.52 \times 10^{+12}$ | 0.1 | $2.49 \times 10^{+10}$ |
| 0.5 | 257 | 10 | $6.73 \times 10^{+5}$ | 5 | $1.19 \times 10^{+17}$ | 75 | $3.66 \times 10^{+10}$ | 0.15 | $1.86 \times 10^{+10}$ |
| 1 | 396 | 20 | $8.24 \times 10^{+5}$ | 10 | $1.13 \times 10^{+17}$ | 100 | $5.04 \times 10^{+7}$ | 0.2 | $1.78 \times 10^{+10}$ |
| 1.5 | 831 | 25 | $1.07 \times 10^{+6}$ | 20 | $1.90 \times 10^{+17}$ | 150 | $4.09 \times 10^{+3}$ | 0.25 | $1.50 \times 10^{+10}$ |
| 2 | $2.47 \times 10^{+3}$ | 30 | $1.75 \times 10^{+6}$ | 30 | $4.51 \times 10^{+17}$ | 200 | $1.57 \times 10^{+5}$ | 0.5 | $7.23 \times 10^{+9}$ |
| 3 | $5.78 \times 10^{+3}$ | 40 | $2.66 \times 10^{+6}$ | 40 | $6.96 \times 10^{+17}$ | 250 | $1.32 \times 10^{+4}$ | 1 | $6.27 \times 10^{+9}$ |
| $m_0 = 110 \mu\text{M}$<br>$n_c = 7.84 \times 10^{-6}$<br>$k+k_n = 6.98 \times 10^{-5}$<br>$(\text{conc}^{-n_2-1} \text{ time}^{-2})$<br>$n_2 = 0.00166$<br>MRE:0.00754 | | $m_0 = 96 \mu\text{M}$<br>$n_c = 0.600$<br>$k+k_n = 257$<br>$(\text{conc}^{-n_2-1} \text{ time}^{-2})$<br>$n_2 = 0.123$<br>MRE:0.00570 | | $m_0 = 55 \mu\text{M}$<br>$n_c = 3.92$<br>$k+k_2 = 4.23 \times 10^{+3}$<br>$(\text{conc}^{-n_2-1} \text{ time}^{-2})$<br>$n_2 = 0.200$<br>MRE:0.00423 | | $m_0 = 55 \mu\text{M}$<br>$n_c = 3.92$<br>$n_2 = 0.2$<br>Global fit:<br>$k+k_2 = 6.71 \times 10^{+5}$<br>$(\text{conc}^{-n_2-1} \text{ time}^{-2})$<br>MRE:0.00905 | | $m_0 = 5.61 \mu\text{M}$<br>$n_c = 0.572$<br>$n_2 = 1.00$<br>$k+k_2 = 3.05 \times 10^{+4}$<br>$(\text{conc}^{-n_2-1} \text{ time}^{-2})$<br>Global fit:<br>$k+k_n = 0.0483$<br>$(\text{conc}^{-n_2-1} \text{ time}^{-2})$<br>MRE:0.0117 | |

**PSMβ1**-heparin interactions follow two different modes below and above 50 µg/mL heparin and are accordingly fitted as two separate sets of data in Amylofit. Data with 0-40 µg/mL heparin are best fitted when k_+_k_n_ values are allowed to vary while keeping the other kinetic parameters n_c_, n_2_ and k_2_k_+_ constant (Fig. S2C). The latter indicates that k_+_ can be considered constant so that heparin mainly affects k_n,_ *i.e.* primary nucleation. The value for k_n_k_+_ remains constant up to ca. 10 µg/mL heparin, after which it increases in a semi-log linear manner (Fig. 2C). Data with 50-250 µg/mL heparin can be fitted by allowing k_n_k_+_ to vary with [heparin] while maintaining a single (global) fit value for k_+_k_2_ (Fig. S2D). The values of k_n_k_+_ decrease dramatically with [heparin], which we again attribute to a reduction in *k*_n_ since *k*_+_ is constant. This time, linearity is only seen when data are plotted in a log-log plot (Fig. 2D). Such log-log linearity is also seen for *e.g.* ligand-binding or protonation/deprotonation systems, *i.e.* strong and specific interactions.

In the absence of heparin, **PSMβ2** follows a primary nucleation and elongation dominated mechanism [11]. However, in the presence of heparin, the best fit is obtained using a secondary nucleation dominated aggregation model where k_+_k_n_ is kept a global constant and k_+_k_2_ is allowed to vary. As with PSMβ2, linearity is only seen when data are plotted in a log-log plot (Fig. 2E). Heparin hence induces secondary nucleation in PSMβ2 while decreasing the aggregation kinetics by inhibiting the peptide’s primary nucleation process (Fig. S2E and Table 1). This is also reflected in the value of k_n_k_+_ which is 3 orders of magnitude lower in the presence of heparin (0.0483 M^-nc^h^-2^) than its absence (48.8 M^-nc^h^-2^).

While δ-toxin on its own did not show any ThT fluorescence increase, heparin dramatically increased its ThT intensity. Since we did not have parameters for heparin-free aggregation, we fitted the data satisfactorily using a secondary nucleation dominated aggregation mechanism with k_n_k_+_ as global fit and k_+_k_2_ as individual fits for each heparin concentration (Fig. S2F). k_+_k_2_ increases with increasing heparin but in a poorly linear manner (Fig. 2F).

It should be noted that fits with better MRE values can be obtained for the kinetic data for PSM peptides in the presence of heparin if the kinetic paramters are set to global fit instead of held as global constants (Fig. S3 and Table S1). This is to be expected since this allows more degrees of freedom during the data fitting. However we find it most consistent to maintain parameters obtained from our previous experiments.

### Heparin has modest effects on the secondary structure and thermal stability of PSM fibrils

Next we addressed whether heparin affected the secondary structures of the fibrillar aggregates. For this we turned to Synchrotron Radiation Circular Dichroism (SRCD) and attenuated total internal reflection Fourier transform infrared (ATR-FTIR) spectroscopy. Each CD and FTIR spectrum was deconvoluted using the DichroWeb server [25, 26] and the OPUS 5.5 software, respectively. Individual CD spectra are shown in Fig. 3A and Fig. S4ABDE with deconvolutions in Fig. 3B and Fig. S4CF; FTIR spectra are presented in Fig. 3C and Fig. S5 with deconvolution results in Fig. 3D. In the absence of heparin, most fibrillar aggregates displayed a single minimum typical of β-sheets as seen for PSMα1 (218nm), PSMα4 (218 nm), PSMβ1 and PSMβ2 (220 nm). These peak positions are in good agreement with previous findings [36]. The FTIR spectra of fibrils related to these four peptides were found to be very similar, with a well-defined intense peak at ∼1625 cm^-1^ indicative of amyloid β-sheet and a minor shoulder at ∼1655 cm^-1^ indicative of α-helical conformation (Fig. 3A and 3C).

**Figure 3:**
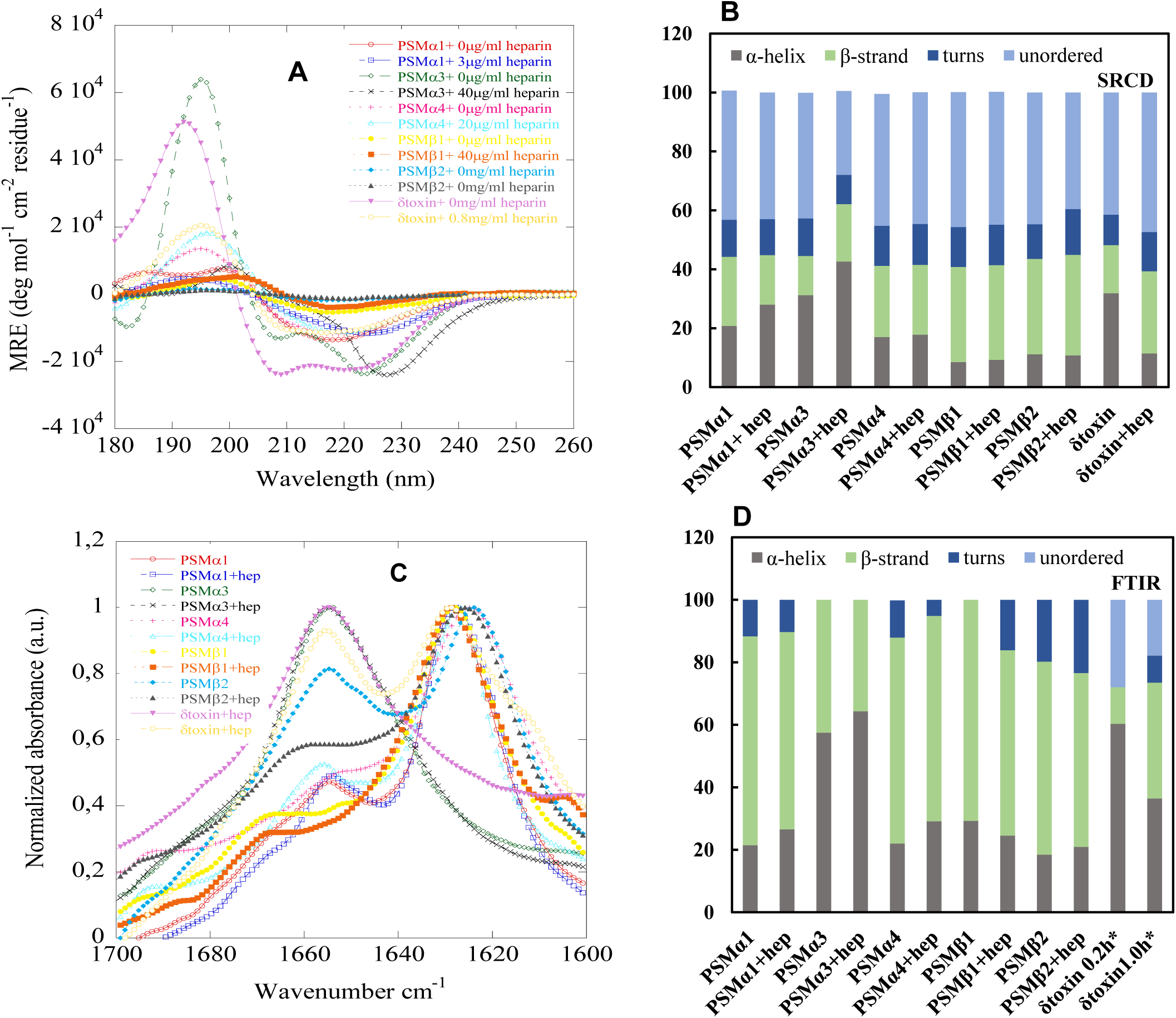
Structural comparison of fibrils formed by different PSMs in the absence and presence of heparin. (A) Synchrotron radiation (SR) Far UV-CD spectra of all PSM fibrils incubated with or without heparin. Fibrillated samples were centrifuged (13k rpm for 30 min), supernatant discarded, and the pellet resuspended in the same volume of milliQ water, (B) Deconvolution of the SRCD spectra from panel A, (C) FTIR spectroscopy of the amide I’ region (1600-1700 cm^-1^) of PSM fibrils formed in the absence and presence of heparin. PSMα1, PSMα4, PSMβ1 and PSMβ2 show a peak at 1625 cm^-1^ corresponding to rigid amyloid fibrils. In contrast, PSMα3 and δ-toxin show main peaks at and 1655 cm^-1^, with the latter indicating more disordered fibrils, (D) Deconvolution of the FTIR spectra from panel C.

Addition of heparin had a variety of effects on fibril structure. PSMα1 showed a slightly shifted minimum ellipticity of 3 and 6 nm in the presence of 1 µg/mL and 3 µg/mL heparin respectively (Fig. S4A). PSMα3 aggregates in presence and absence of heparin show cross-α-helical structure in both SRCD and FTIR spectra, in good agreement with previous reports [37, 38]. The SRCD and FTIR spectra for PSMα4 in absence and presence of heparin were internally consistent (Fig. 3AC and Fig. S4B). An FTIR peak around ∼1690 cm^-1^ suggests anti-parallel beta-sheets. SRCD spectra of PSMβ1 and PSMβ2 heparin-free aggregates are similar to those in the presence of heparin (Fig. 3A and Fig. 3B, S4D). Further, SRCD and FTIR spectra clearly indicate that heparin favors amyloid formation in δ-toxin and significant secondary structural changes were observed in the presence of various concentrations of heparin (Fig. 3A and Fig. S4EF).

The thermal stability of the PSM aggregates in absence and presence of heparin was evaluated by CD spectroscopy. Fig. S6 shows thermal scans monitored by ellipticity at 220 nm for fibrils (PSMα1, PSMα3 and PSMα4) from 25° C to 95° C. Neither PSMα1 and PSMα4 nor βPSMs fibrils undergo any significant loss in signal up to 95°C, indicating thermally stable β-sheet structure (Fig. S6A, B). However, PSMα3 fibrils are thermally unstable both conditions in the absence and presence of heparin, as a loss of structure is seen above 50°C (Fig. S6A). Hence heparin does not change the thermal stability of the fibrils.

### Heparin shows variable effects on the fibril morphology of PSMs

The morphologies of the peptide fibrils in the absence and presence of heparin were analyzed by transmission electron microscopy (TEM). TEM analysis of PSMα1 incubated in the absence and presence of heparin showed amyloid-like fibrils in both cases (Fig. 4A and 4B) though fibrils prepared with heparin were slightly thicker. No aggregated species were observed for PSMα2 (Fig. S7A-B) in both conditions, consistent with the lack of increase in ThT fluorescence upon incubation with and without heparin. In the absence of heparin, PSMα3 formed long rod like fibrils (Fig. 4C) which in the presence of heparin shows very similar fibrillar structure (Fig. 4D).

**Figure 4:**
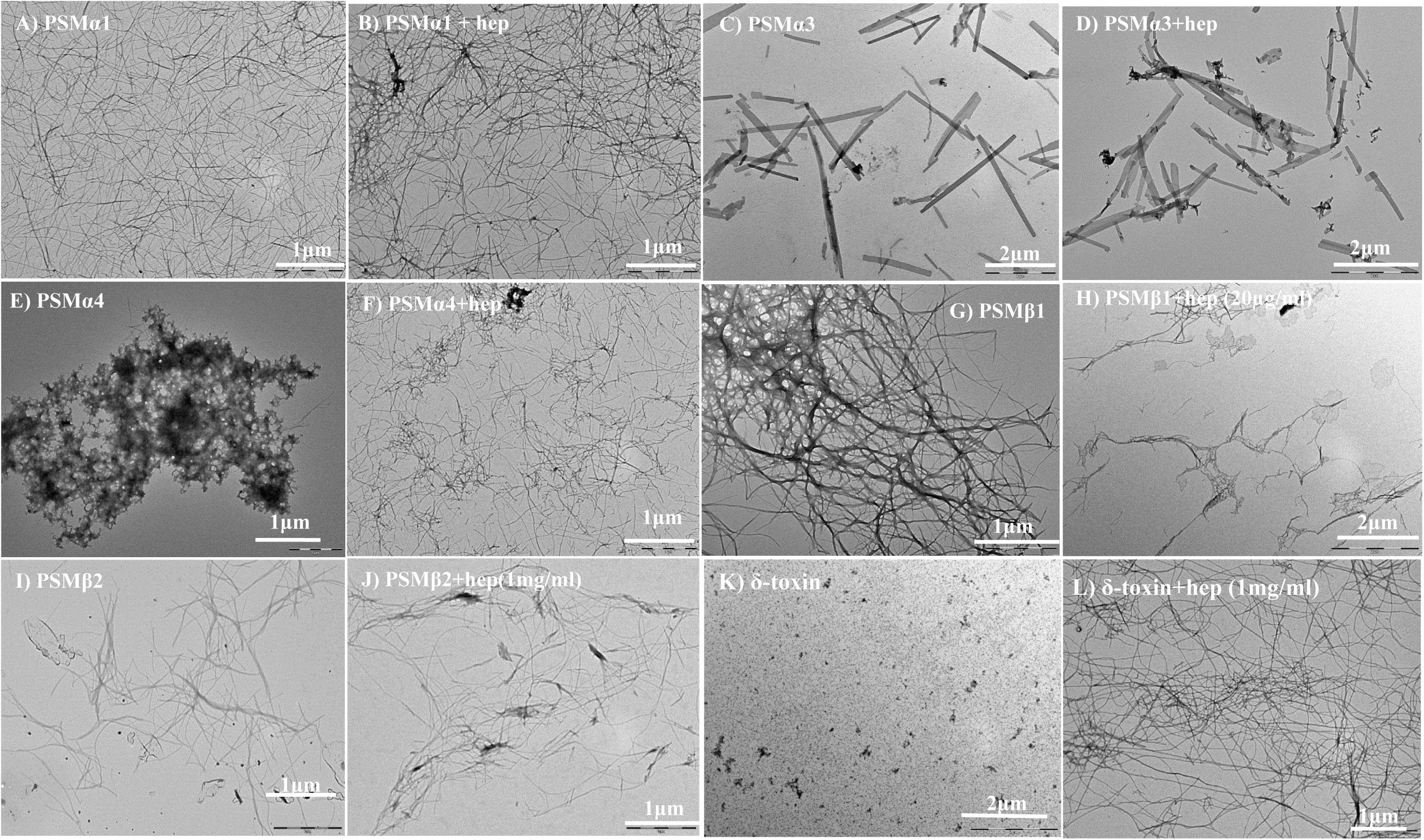
Electron microscope images of fibrils formed from PSMs in the absence and presence of heparin. TEM micrographs of (A) PSMα1 without heparin, (B) PSMα1 with 3µg/mL heparin, (C) PSMα3 without heparin, (D) PSMα3 with 40 µg/mL heparin, (E) PSMα4 without heparin, (F) PSMα4 with 20 µg/mL heparin, (G) PSMβ1 without heparin, (H) PSMβ1 with 40 µg/mL heparin (I) PSMβ2 without heparin, (J) PSMβ2 with 1 mg/ml heparin (K) δ-toxin without heparin and (L) δ-toxin with 1 mg/ml heparin. Note that scale bars vary between panels.

Heparin also encouraged formation of PSMα4 fibrils (Fig. 4E, F). On their own, PSMα4 displays very thin fibrils visible at higher magnification with some distribution of spherical aggregates organized into small clusters on the grid which upon incubation with heparin show nicely separated fibrillar structure (Fig. 4E). PSMβ1 on its own formed highly ordered arrays of laterally associated fibers (Fig. 4G) which with heparin changed to disentangled thin aggregates (Fig. 4H and Fig. S7C). We do not observe significant morphological difference between PSMβ2 fibrils obtained in the absence and presence of heparin (Fig. 4I and 4J). Finally, TEM of δ-toxin confirmed the aggregation potential of heparin. While there were no visible fibrils in δ-toxin on its own, heparin led to large networks of thin fibers (Fig. 4K and 4L, Fig. S7D).

### Interactions between heparin and PSM residues analyzed by peptide arrays

We explored the interaction between fluorescein-labeled heparin and PSM sequences using a peptide array chip displaying 10-residue immobilized peptides in staggered arrangements (Tables S3 and S4) . Thanks to the fluorescein label, it was possible to quantitate the amount of heparin bound to each peptide fragment (Fig. 5). All α-type PSMs bind more heparin than βPSM, but the intensity is not equally distributed along the length of each PSM. High heparin affinity is shown by peptides corresponding to the N-terminal half of α-PSMs and the middle and C-terminal part of β-PSMs. We attempted to probe possible correlations between signal intensity and the peptides’ physical-chemical characteristics such as charge and hydrophobicity [39]. As shown in **Table 2**, the sequence charge is the most significant contributor, especially for βPSMs, with higher positive charge leading to higher signal intensities. This is to be expected in view of heparin’s highly anionic nature.

**Figure 5:**
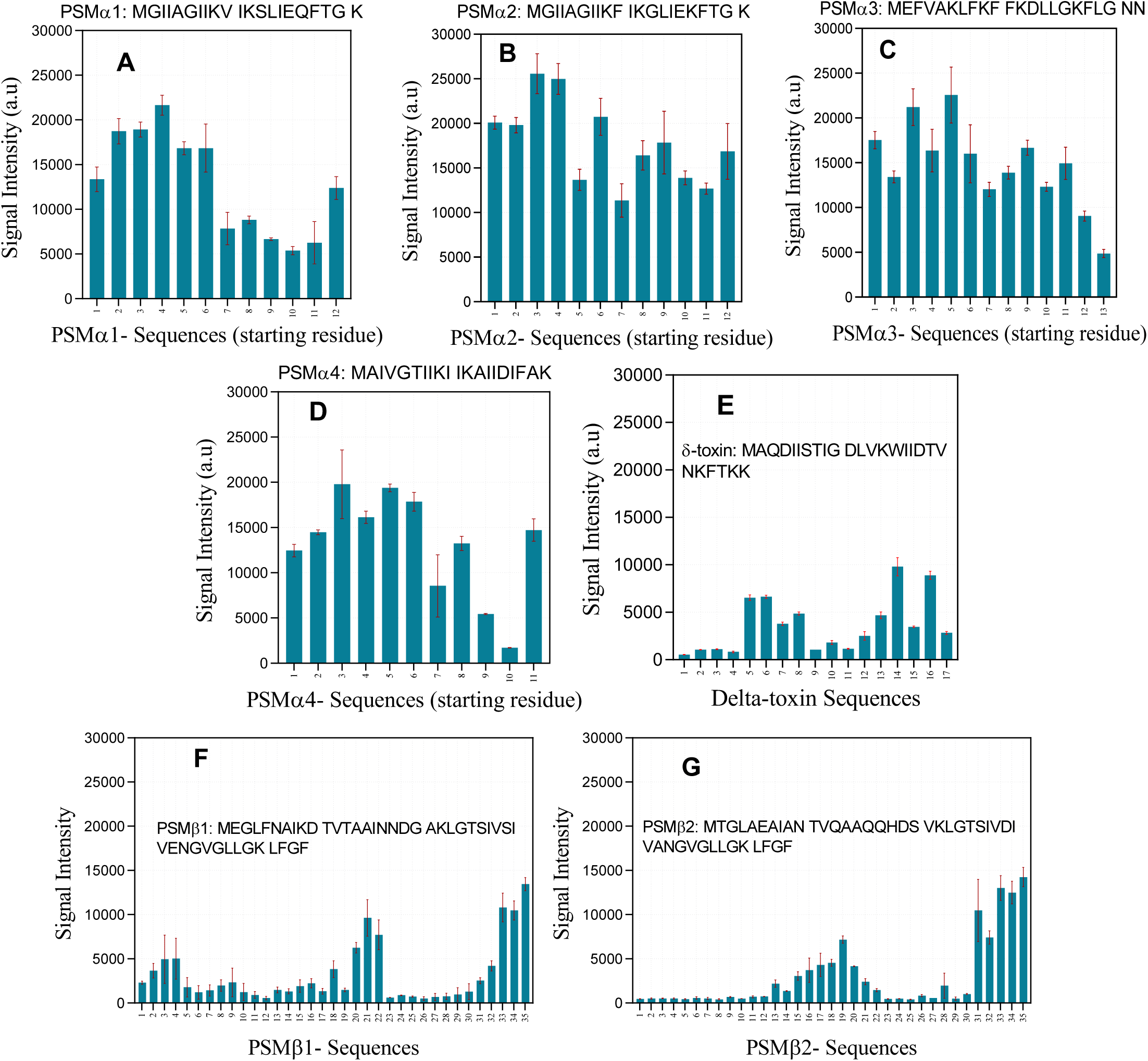
Interaction of fluorescein-labeled heparin with different PSM peptides displayed on a peptide array. Data provides signal intensity from different PSM sequences interacting with heparin. Full PSM sequences are provided in each panel. (A) PSMα1, (B) PSMα2, (C) PSMα3, (D) PSMα4, (E) δ-toxin, (F) PSMβ1, (G) PSMβ2. For each spot, the number on the x-axis gives the residue position in the intact PSM sequence, corresponding to the starting residue in the spot’s 10-mer peptide.

**Table 2.**
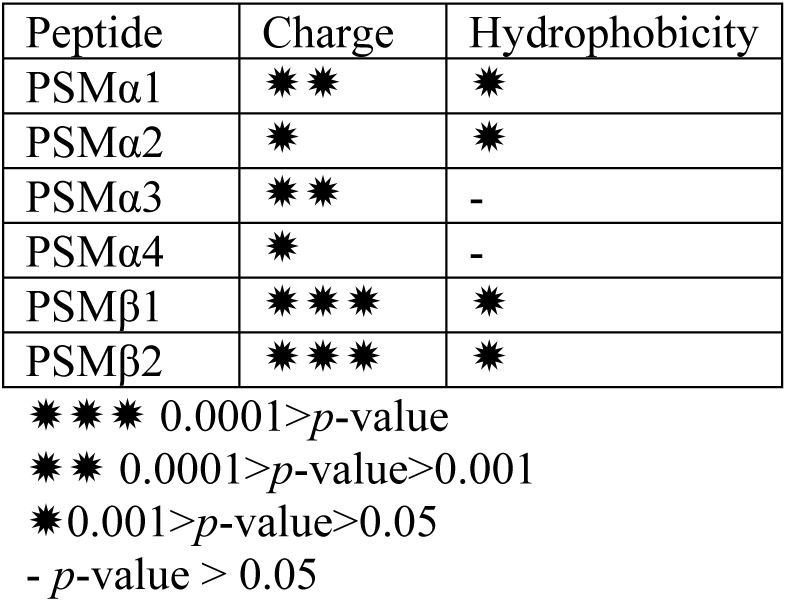
*p*-values from multiple regression analysis of the correlation between signal intensity and the two variants charge and hydrophobicity.

Note however that *a priori* we cannot predict whether high binding by heparin would promote aggregation (*e.g.* by forming a template for the extended state, leading to amyloid) or inhibit it (by sequestering monomers from interacting with other monomers). According to our ThT-assays (Fig. 1) heparin accelerates the fibrillation of PSMα1, PSMα3, PSMα4, δ-toxin and PSMβ1 at low heparin concentrations, whereas it inhibited PSMβ1at high heparin concentrations as well as PSMβ2. We conclude from this that the high affinity of heparin to α-PSMs’ N-termini promotes fibrillation, whereas binding to the middle and C-terminal regions of βPSMs inhibits fibrillation.

To elucidate the role of individual residues in the heparin interaction, we carried out an Ala scan of all PSMs (Fig. 6).

**Figure 6:**
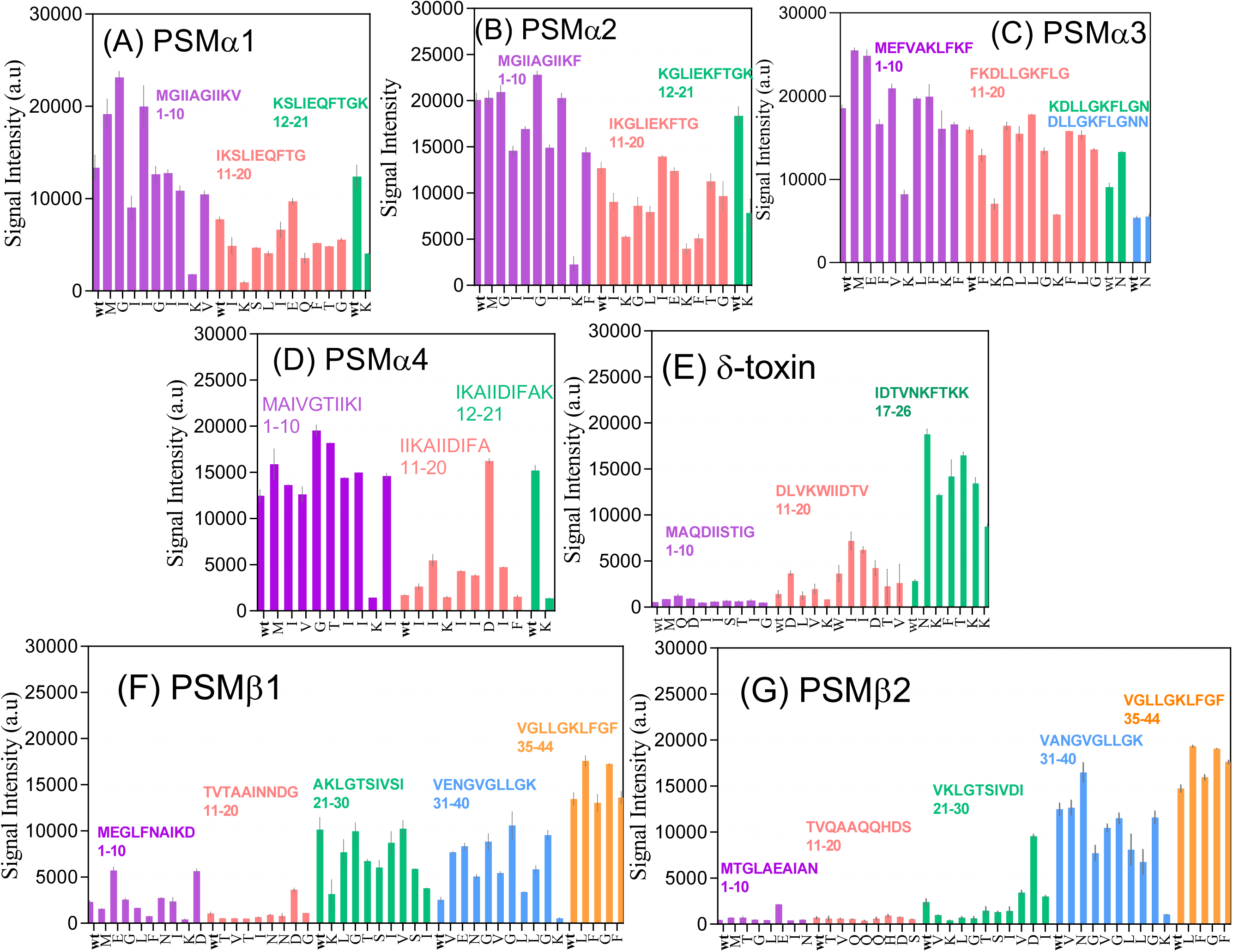
Interaction of fluorescein-labeled heparin with peptide-array peptide sequences designed for Ala scans of different PSMs. Data provides signal intensity from different Ala-scanned PSM sequences interacting with heparin. Each PSM peptide was divided into 10-residue sequences, each with their own color; in each peptide, positions were individually replaced by Ala from left to right. “wt” is the initial peptide before starting Ala scan. Each letter on the x-axis show the residue that is replaced by Ala, (A) PSMα1, (B) PSMα2, (C) PSMα3, (D) PSMα4, (E) δ-toxin, (F) PSMβ1, (G) PSMβ2.

In PSMα1, an *increase* in signal intensity compared to wildtype was caused by mutations M1A, G2A and I4A (which maintain the same charge but lead to either a fall or a rise in hydrophobicity) (Table S1). A *decrease* in signal was caused by removal of positive charge (K9A, K12A, and K21A). This illustrates clearly how electrostatic interactions are the main (but not the only) drivers of heparin-peptide interactions (Fig. 6A).

Similarly, K-to-A mutations in PSMα2 and also in the rest of PSM peptides (including PSMα3, PSMα4, δ-toxin, PSMβ1 and PSMβ2) decreased heparin binding, but so did loss of hydrophobicity: mutations I3A, I6A and F18A for PSMα2 and I3A, I11A and L14A for PSMα1 led to a 25-50% drop in intensity. We were unable to obtain ThT fibrillation curves for PSMα2, thus we cannot conclude what effect binding would have on its fibrillation kinetics.

Moreover, removal of negative charge (e.g. for E and D to A residues) leads to an increase in signal intensity, but mutations altering hydrophobicity had an even more marked increase in binding for peptides with same charges (Table S2). Mutation M1A also led to increase in signal for PSMα3 and PSMα4 same as PSMα1. For PSMα3, K-to-A mutations led to the highest decrease in signal intensity, again emphasizing the role of charge. For PSMα4 except the mutations affect the charge, the truncation mutations T6A, I12A, I14A, I15A and I17A increased binding as did the insertion mutation G5A. G-to-A/T mutations increase hydrophobicity while I-to-A decrease it; neither affect charge. Interestingly, the I-to-A mutation led to a drop in signal for PSMα1 (I4A, I11A) and PSMα2 (I3A, I4A, I6A and I11A) but increased it for PSMα4 and δ-toxin (I16A and I17A), indicating different roles for Ile.

For δ-toxin, the Ala scan of first 10 residues did not show major changes. For the second 10 residues, almost all mutations led to higher signal intensity and the most important mutations are D11A, W15A, I16A, I17A and D18A. Ala scans of the last 6 residues increased the signal intensities to the highest level. Remarkably, this was also the case for 3 K-to-A mutations. Note that these 6 spots are the only positively charged spots in Ala scan of δ-toxin (which does not fibrillate in the absence of heparin), and they could be binding partners for heparin that promote fibril formation. Also in PSMβ1 and PSMβ2, removal of anionic Glu/Asp led to increased binding, while removal of Lys decreased signal intensity (Fig. 6 F and G).

**Table 3** shows that for PSMβ1 and PSMβ2, charge is most strongly correlated with binding (*p*-values 6.12*10^-6^ and 3.01*10^-9^ respectively), while hydrophobicity has a much weaker effect on PSMβ1 (*p*-value 0.025) and PSMβ2 (*p*-value 0.313). Based on multiple regression analysis with two variants charge and hydrophobicity, the predicted signal intensities nicely fit the measured signals, indicating the central importance of these two parameters in the interaction between PSMs and heparin (Fig. S8).

**Table 3.**
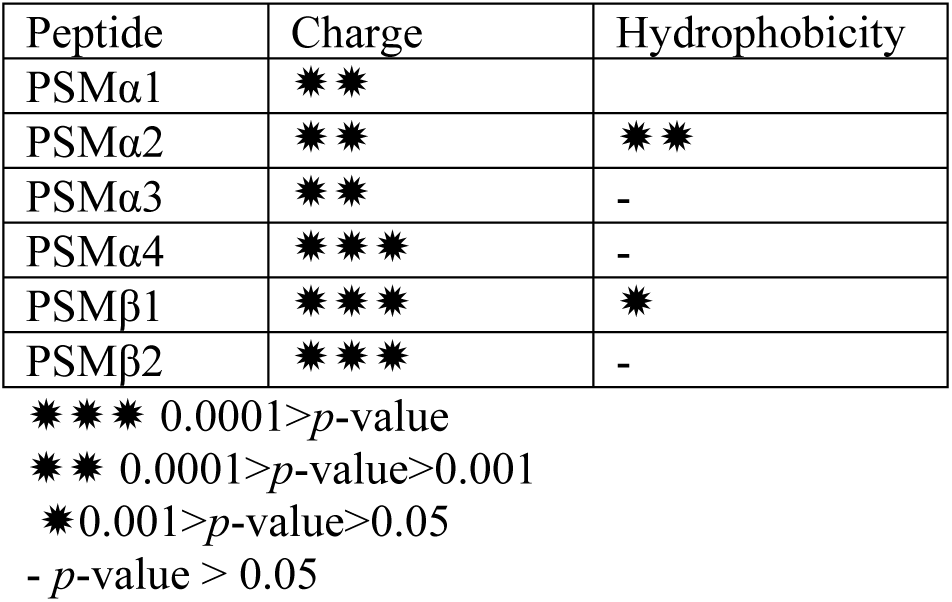
*p*-values from multiple regression analysis of the correlation between signal intensity and the two variants charge and hydrophobicity

### Heparin promotes biofilm formation both in the presence and absence of PSMα/β

To put our observations in a biological context, we investigated how heparin affects biofilm formation. Accordingly, we incubated *S.aureus* Newman strain as a model of *S.aureus* human infections having a robust virulence phenotype and ability to form biofilm [40, 41] and three different PSM mutants of this strain with 10-200 µM of heparin. Incubation of wild type *S. aureus* with heparin increased the amount of biofilm significantly at > 10 µg/mL heparin (Fig. 7). A strain that only produces PSMα (ΔPSMβ) show a higher biofilm formation even at low [heparin] compared to ΔPSMα; this could be caused by the ability of heparin to induce PSMα fibrillation. However, an increase in the biofilm formation of ΔPSMα/β in the presence of heparin indicates that other mechanisms are also involved in biofilm formation.

**Figure 7:**
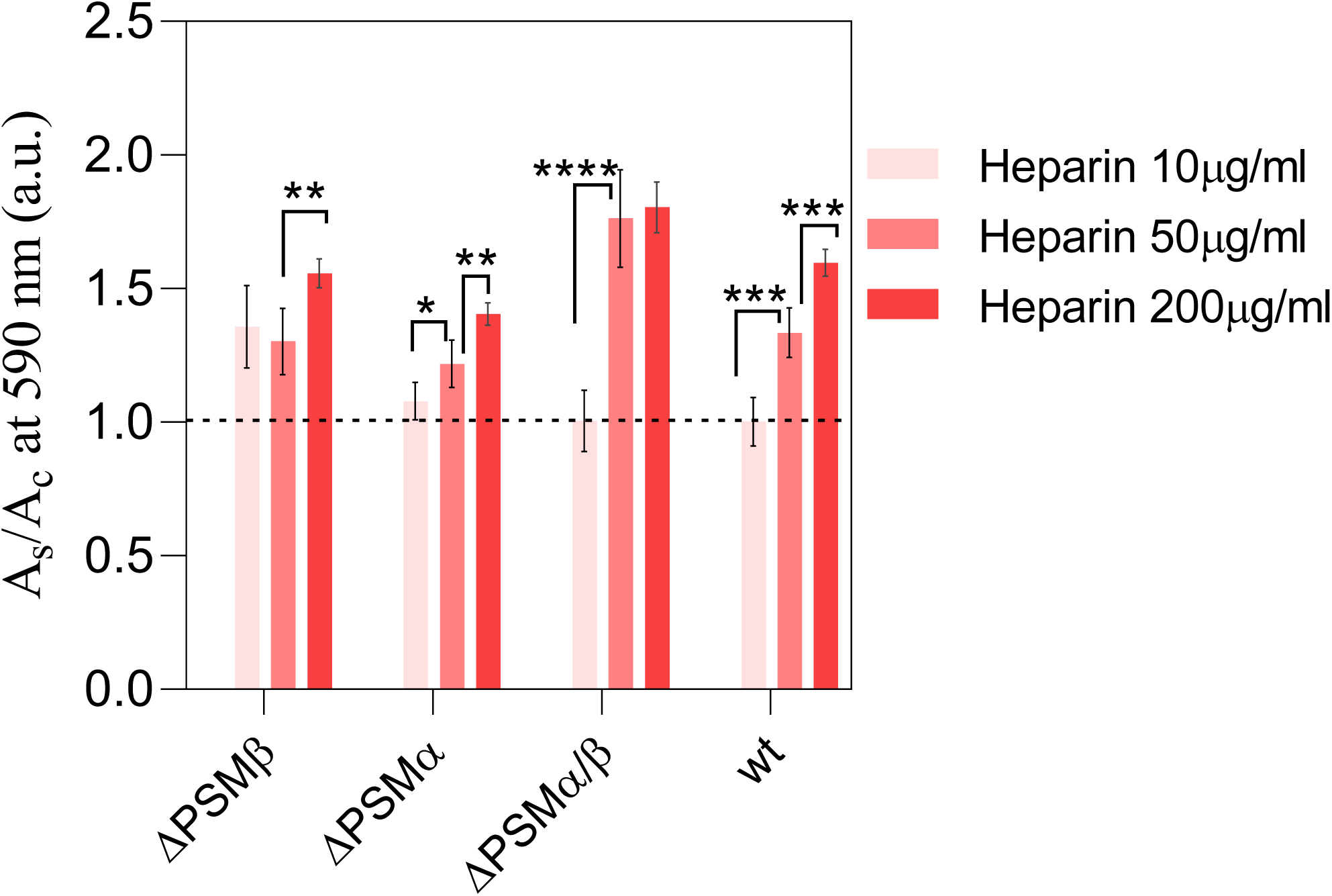
Biofilm formation in the presence of heparin. A_s_/A_c_ is the absorbance of samples at 590 nm (indicative of the amount of biofilm produced) in the presence of heparin divided by absorbance of the control (same strain grown in the absence of heparin). Mean ± SD, n=3, * *p* < 0.05, ** *p* < 0.002, *** *p* < 0.0002, **** *p* < 0.0001.

## Discussion

*Staphylococcus aureus* is an important human pathogen causing many different hospital- and community-associated infections. This is promoted by its ability to form biofilm, aided by amyloid-forming PSMs [42], and thus increase resistance to antibiotics [43]. *S.aureus* is particularly prone to form biofilm on catheters; furthermore heparin is commonly used as anticoagulant in these catheters [17]. This inspired us to investigate the effect of heparin on PSMs fibrillation.

### Heparin shows a range of effects on aggregation which are very sensitive to peptide sequence

Our study highlighted that heparin accelerates fibril formation for αPSMs (PSMα1, PSMα3, PSMα4 and δ-toxin) at concentrations as low as 1μg/mL (PSMα1), though the effect was generally seen in the range 0.02-1 mg/mL. In contrast, heparin inhibits the fibrillation of the two PSMβs. The interaction is mainly driven by electrostatic interactions between PSMs and heparin as seen for many other protein-heparin interactions [44].

In the PSMα family, heparin induces fibrillation by reducing nucleation time and enhancing the ThT end-level. It also induces aggregation of the otherwise non-aggregating δ-toxin. PSMβ1 is inhibited in two different modes: below 50 μg/mL heparin, heparin inhibits the nucleation step but does not affect growth rates, but above 50 μg/mL heparin increased growth rate and ThT end-levels while increasing the lag phase. Heparin had mixed effects on PSMβ2, increasing the lag-time as well as ThT-end levels. It is remarkable that PSMβ1 and PSMβ2 show very distinct aggregation behavior despite their high similarity. PSMβ1 aggregates efficiently at very low concentrations and responds in a bimodal manner to heparin, with different behavior ats at low versus high heparin concentrations. Amylofit analysis reveals that the k_+_k_2_ is mostly affected in the α-PSM peptide whereas incubation with heparin leads to variations in k_+_k_n_ for β-PSM peptides, thus highlighting the effect of heparin on the nucleation phase of aggregation for the β-PSM peptides in particular.

The marked increase in the final ThT-level is also seen as a higher level of fibril formation. Similar effects have been reported for amyloidogenic proteins like α-synuclein and transthyretin [21, 23]. Heparin co-pellets with the aggregates, showing strong binding and a possible templating role in fibrillation [21, 23, 45, 46]. The shortening of the lag phase seen for PSMα1 and PSSMα3 might be caused by heparin’s induction of a conformational changes favoring fibrillation process or by stabilizing early-stage aggregates [14, 23, 47].

### Cationic heparin-binding motifs in the αPSMs family may drive fibrillation

Heparin-binding domains often contain a high proportion of positively charged Lys and Arg which can interact with anionic GAGs [14]. These residues often occur as motifs with basic amino acids in close proximity like XBBBXXBX and XBBXBX sequences where B and X are basic and nonbasic residues respectively [48, 49]. The proximity of B residues likely leads to cooperative binding effects. Such motifs are found in the αPSMs family, *e.g.* “KVIK” in PSMα1, “KLFK” in PSMα3, “KIIK” in PSMα4 and “KFTKK” in δ-toxin. For βPSMs the Lys residues are further from each other and their net charge at pH 7 is overall negative. This might explain the difference in behavior. Our peptide array data also confirm the importance of positively charged residues (mainly Lys) whose replacement by Ala abolished heparin binding. Further, removal of negatively charged residues (D and E residues) increased the binding of heparin. Both confirm the importance of electrostatic interaction between PSMs and heparin. Alanine scan of peptides on peptide array indicated that charge is more important than hydrophobicity as K-to-A mutations increased the hydrophobicity but did not increase the binding of heparin (table S2). The structure of PSMα3 fibrils reveals that some Lys residues are not involved in intermolecular fibrillar contacts but are speculated to be related to the cytotoxicity of the peptide towards human cells (*e.g.* through interactions with the membrane) as mutations of these residures to Ala results in reduced toxicity [50]. This implies that these residues can take part in electrostatic interactions with heparin without interfering with amyloid formation.

Structural analysis of fibrils demonstrate typical β-sheet fibrils for PSMα1 (218nm), PSMα4 (218 nm), PSMβ1 and PSMβ2 (220 nm) as reported [11, 36, 51] and this is largely unaffected by heparin. Simiarly, the characteristic but unusual cross-α fibrils of PSMα3 are maintained in the presence of heparin [9, 50]. Heparin led the non-aggregating δ-toxin peptide (which forms α-helices in solution) to form β-sheet amyloid fibrils. All fibrils show high thermal stability except the cross-α PSMα3 fibrils which are unstable above 50°C. Similar modest thermal stability has been reported for fibrils of PSMα3-LFKFFK segment [51]. Thus the cross-α structure may be inherently less stable than cross-β fibrils due to the difference in the type of intermolecular contacts.

Incubation of *S. aureus* Newman strain and its *psm* mutants in presence of heparin show biofilm formation significantly promoted in wild type and ΔPSMβ, but not so pronounced in the case of ΔPSMα. Gratifyingly, this is consistent with the inducing and inhibiting effect of heparin on fibrillation of αPSMs and βPSMs respectively. The role of complemenmtary electrostatics is also confirmed by the observation that positively charged polysaccharides like chitosan do not increase biofilm formation [17, 52, 53].

In summary, our study uncovered a diversity of mechanistic effects of heparin on the fibrillation of PSMs. There were modest differences in the kinetics of the heparin-stimulated fibrillation reaction of PSMs, with the kinetics being fastest with PSMα3 and slowest with δ-toxin and there are significant differences in the seven PSM peptides’ affinities for heparin. While heparin promotes fibrillation of αPSMs and δ-toxin, it inhibited but did not abolish βPSM fibrillation (Fig. 9), consistent with its ability to promote *S. aureus* biofilm formation. Heparin mostly targets the nucleation step and thus the lag phase, while increasing ThT end levels suggest higher levels of fibrillation. Furthermore, our data demonstrate that positively charged residues close to each other in αPSMs and δ-toxin provide suitable region to stabilize binding of the highly negative charge heparin. Additionally, in contrast to most previous studies showing only the effects of heparin as a promoter of fibrillation [54, 55], our results demonstrated that heparin has a dual effect and it acts as an inducer or inhibitor in the fibrillation of PSMs, which main contribute both to the integrity and dynamics of formation of biofilms.

## Acknowledgments

D.E.O. gratefully acknowledges support by the Independent Research Foundation Denmark | Technical Sciences (grant no. 6111-00241B) and the Independent Research Foundation Denmark | Natural Sciences (grant no. 8021-00208B). M.A. gratefully acknowledges support by Aarhus University Research Foundation. Furthermore, we acknowledge the award of beam time on the AU-CD beam line at ASTRID2, under project number ISA-20-1013 and Dr. Nykola Jones for assistance for data collection.

## Author contribution

Z.N., M.Z., M.A. and D.E.O. designed experiments, M.Z. and Z.N. carried out the experiments, M.Z., Z.N., M.A. and D.E.O. analyzed the data, Z.N., M.Z., M.A. and D.E.O. wrote, reviewed and edited the manuscript while D.E.O. and M.A. acquired funding.

---

**Figure S1:**
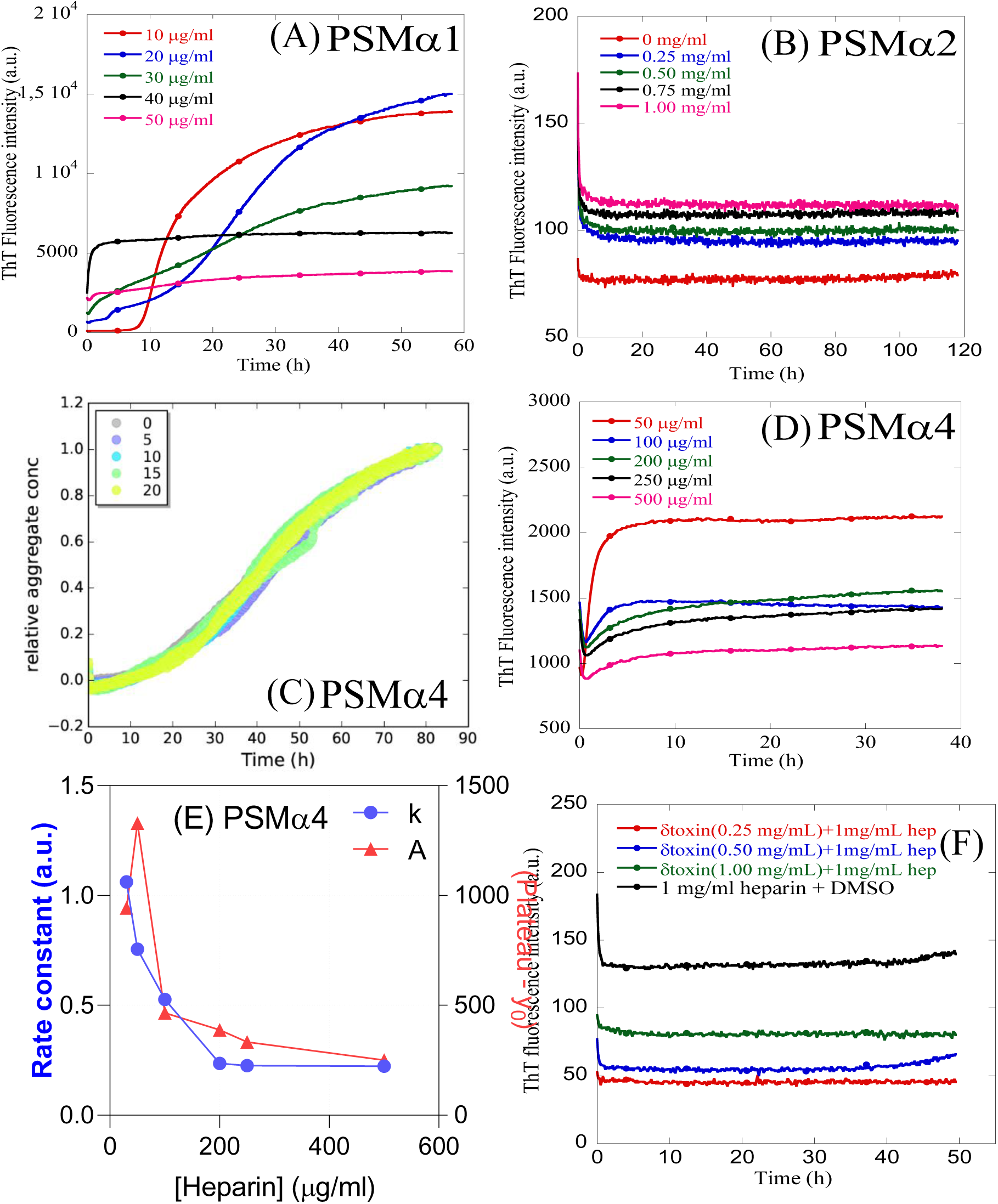
Effect of heparin concentration on PSMs fibrillation kinetics under quiescent conditions. (A) 0.25 mg/mL PSMα1 with 10-50 µg/mL heparin. (B) 0.25 mg/mL PSMα2 with 0-1 mg/mL heparin. (C) Normalized ThT-curve for PSMα4 incubated with 0-20 µg/ml heparin. (D) 0.25 mg/mL PSMα4 with 50-250 µg/mL heparin. (E) Kinetic parameters for PSMα4 versus heparin concentration. Data for 30-500 µg/mL heparin in panel D were fitted with an exponential decay to yield rate constant *k* and amplitude *A*. (F) 0-1 mg/ml monomeric δ-toxin in presence of 1 mg/ml heparin. Data are representatives of triplicate experiments.

**Figure S2:**
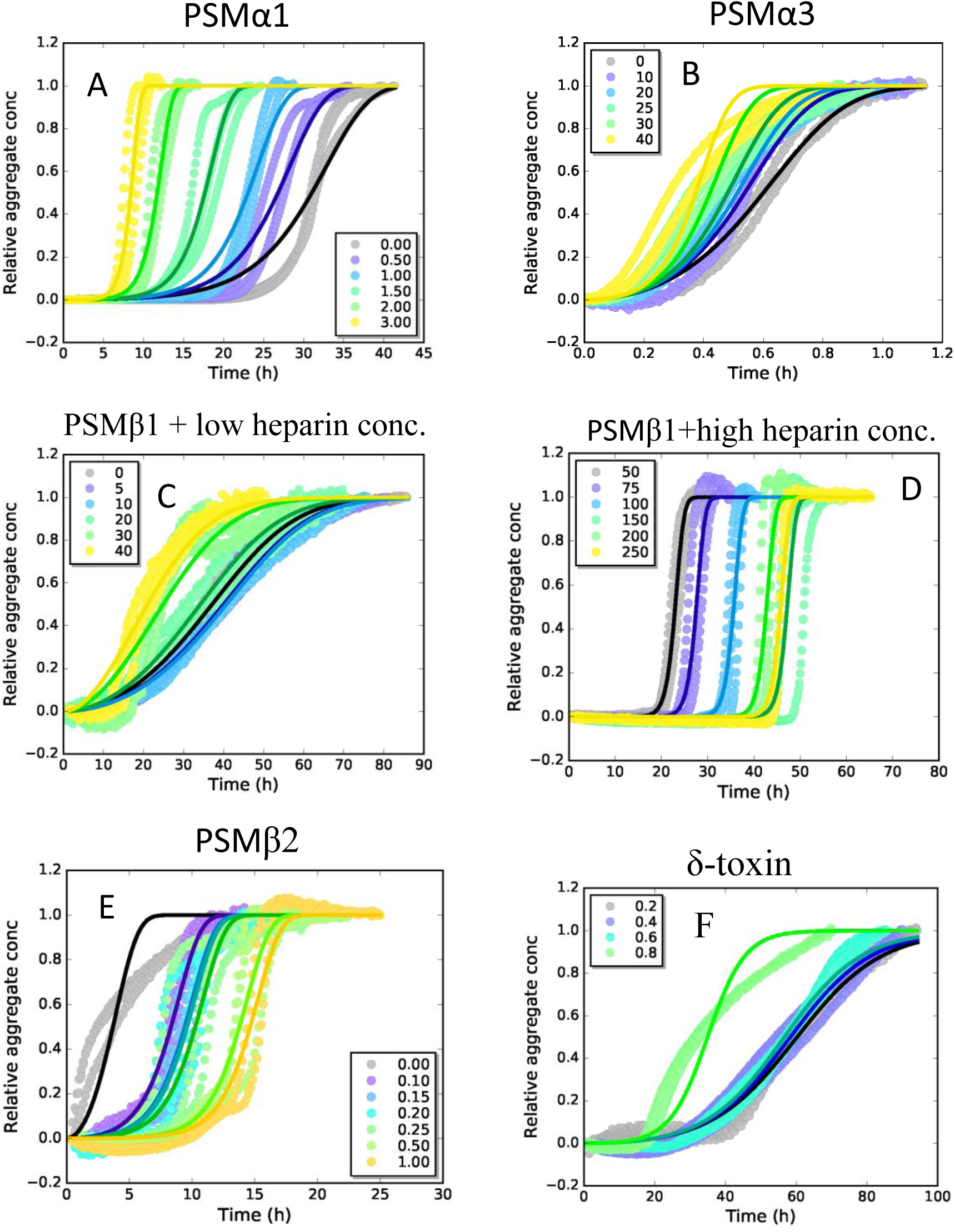
Fitting of aggregation kinetic data for PSM peptides in the presence of heparin using Amylofit. The heparin concentration in µg/mL is indicated for each curve, all data fitted to a secondary nucleation dominated model using global constants: n_c_,n_2_, k_+_k_n_, individual fit: k_+_k_2_ (for PSMβ1: individual fit: k_+_k_n_ and global constant k_+_k_n_). (A) Fitting of PSMα1 kinetic data at 0.25 mg/mL PSMα1 in the presence of 0-3 µg/mL heparin. (B) Fitting of PSMα3 kinetic data at 0.25 mg/mL PSMα3 in the presence of 0-40 µg/mL heparin. (C) Fitting of PSMβ1 kinetic data at 0.025 mg/mL PSMβ1 in the presence of 0-40 µg/mL heparin. (D) Fitting of PSMβ1 kinetic data at 0.025 mg/mL PSMβ1 in the presence of 50-250 µg/mL heparin. (E) Fitting of PSMβ2 kinetic data at 0.25 mg/mL PSMβ2 in the presence of 0.1-1 µg/mL heparin. (F) Fitting of δ-toxin kinetic data at 0.3 mg/mL δ-toxin in the presence of 0.2-0.8 µg/mL heparin fitted to a secondary nucleation dominated model using global constants: n_c_,n_2_, k_+_k_n_, individual fit: k_+_k_2_.

**Figure S3:**
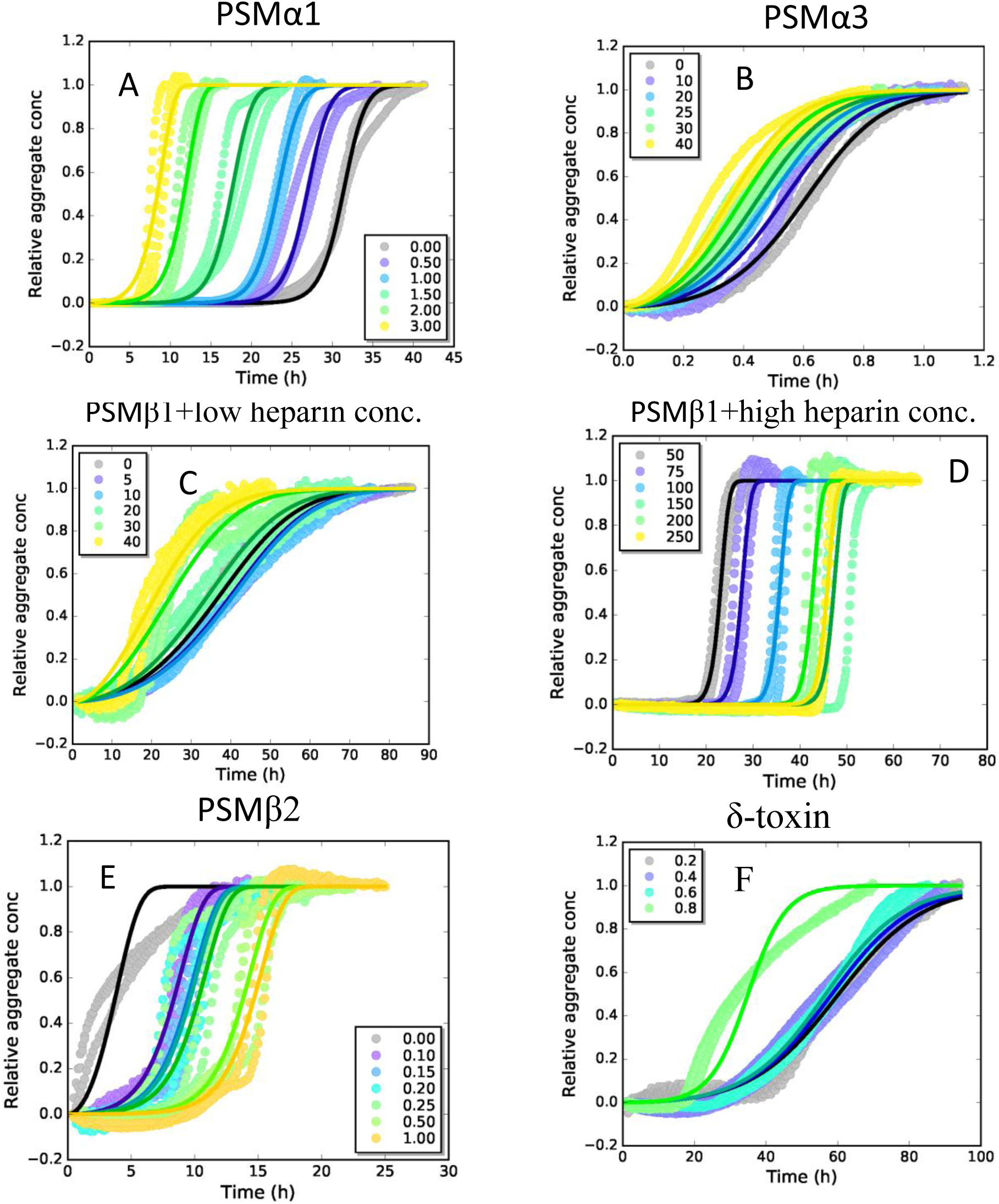
Fitting of aggregation kinetic data for PSM peptides in the presence of heparin. The heparin concentration in µg/mL is indicated for each curve, all data fitted to a secondary nucleation dominated model using global fitting of: n_c_, n_2_, k_+_k_2_, individual fit: k_+_k_n_. A) Fitting of PSMα1 kinetic data at 0.25 mg/mL PSMα1 in the presence of 0-3 µg/mL heparin. B) Fitting of PSMα3 kinetic data at 0.25 mg/mL PSMα3 in the presence of 0-40 µg/mL heparin. C) Fitting of PSMβ1 kinetic data at 0.025 mg/mL PSMβ1 in the presence of 0-40 µg/mL heparin. D) Fitting of PSMβ1 kinetic data at 0.025 mg/mL PSMβ1 in the presence of 50-250 µg/mL heparin. E) Fitting of PSMβ2 kinetic data at 0.25 mg/mL PSMβ2 in the presence of 0.1-1 µg/mL heparin. F) Fitting of δ-toxin kinetic data at 0.3 mg/mL δ-toxin in the presence of 0.2-0.8 µg/mL heparin fitted to a secondary nucleation dominated model using global constants: n_c_,n_2_, k_+_k_n_, individual fit: k_+_k_2_.

**Figure S4:**
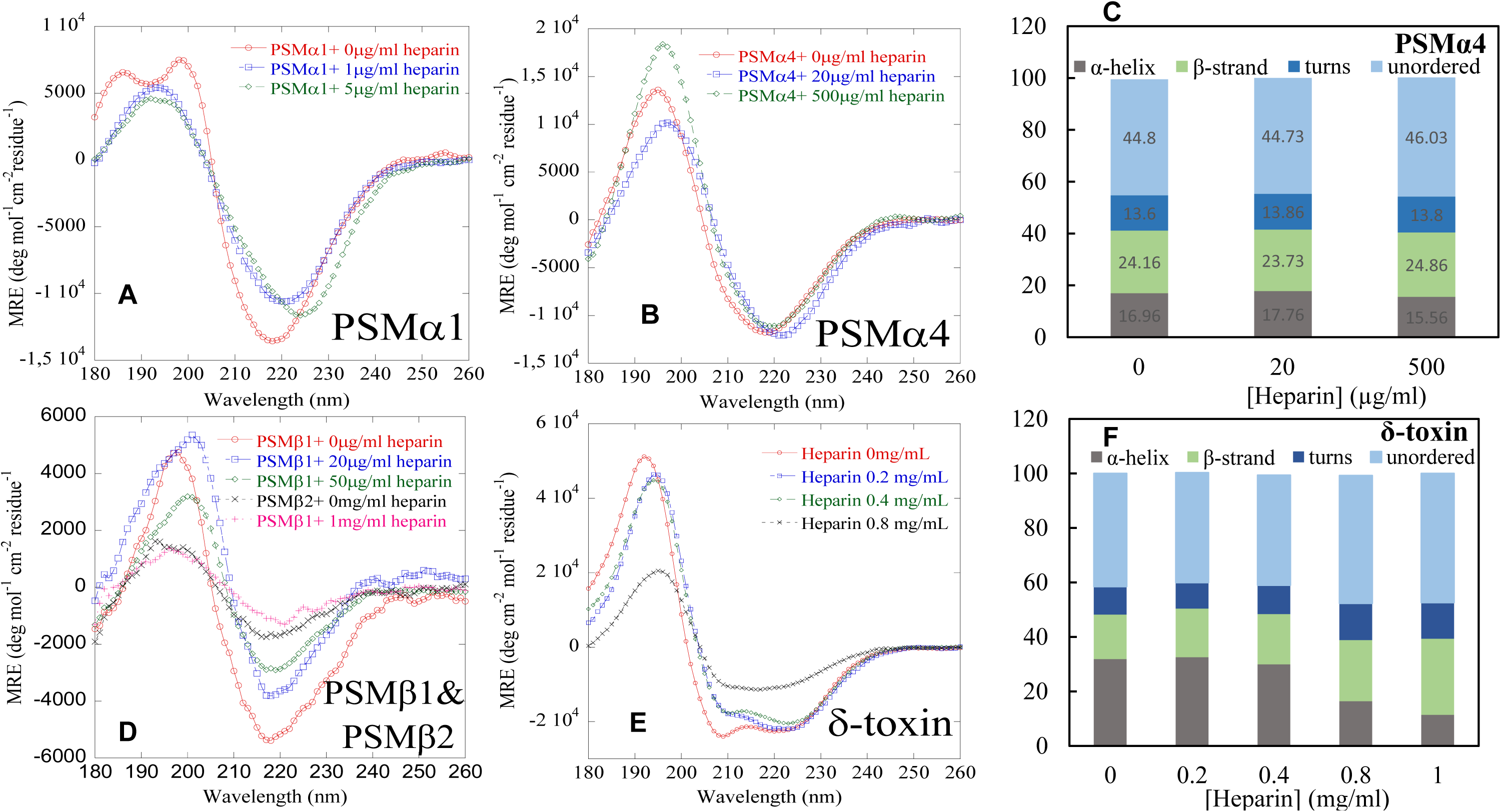
Conformational conversion of PSMs followed by far-UV CD. (A) SRCD spectra of PSMα1 (0.25 mg/mL) in absence and presence of different concertation of heparin. (B) SRCD spectra of PSMα4 (0.25 mg/mL) in absence and presence of different concertation of heparin. (C) Deconvolution of the SRCD spectra of fibrils of PSMα4 (with/out heparin) into the individual structural components. (D) Synchrotron radiation (SR) Far UV-CD spectra of all β-group of PSMs fibrils incubated with/out heparin. (E) Synchrotron radiation (SR) Far UV-CD spectra of δ-toxin fibrils incubated in absence and presence of different concentrations of heparin. (F) Deconvolution of the SRCD spectra of fibrils of δ-toxin (with/out heparin) into the individual structural components.

**Figure S5:**
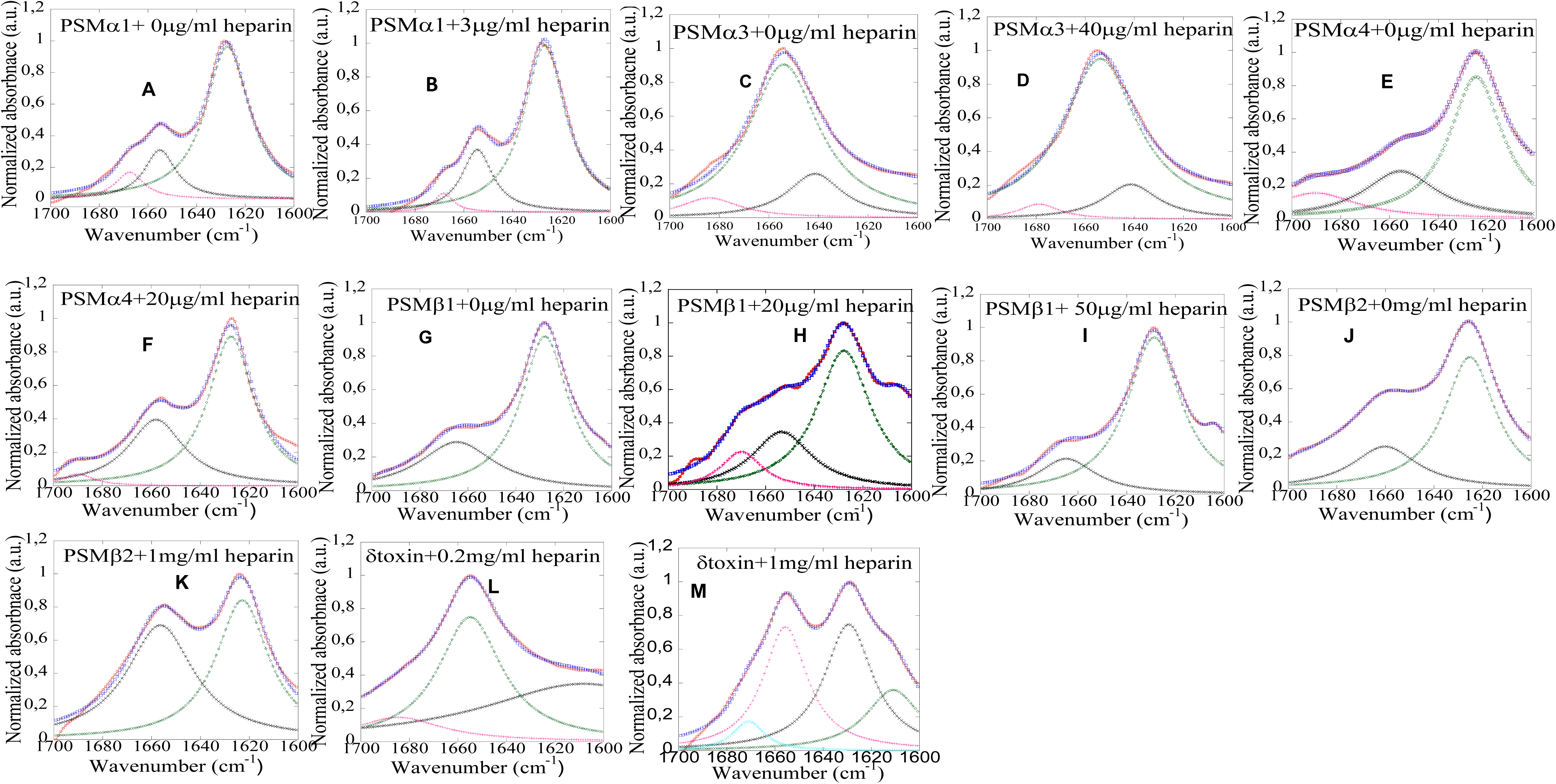
FTIR analysis of the secondary structure of PSMs peptides in absence and presence of heparin. The data were processed by baseline correction and interfering signals from H_2_O and CO_2_ were removed using the atmospheric compensation filter. Peak positions were assigned where the second order derivative had local minima and the intensity was modeled by Gaussian curve fitting using the OPUS 5.5 software. Each panel represent FTIR spectra and second order derivative.

**Figure S6:**
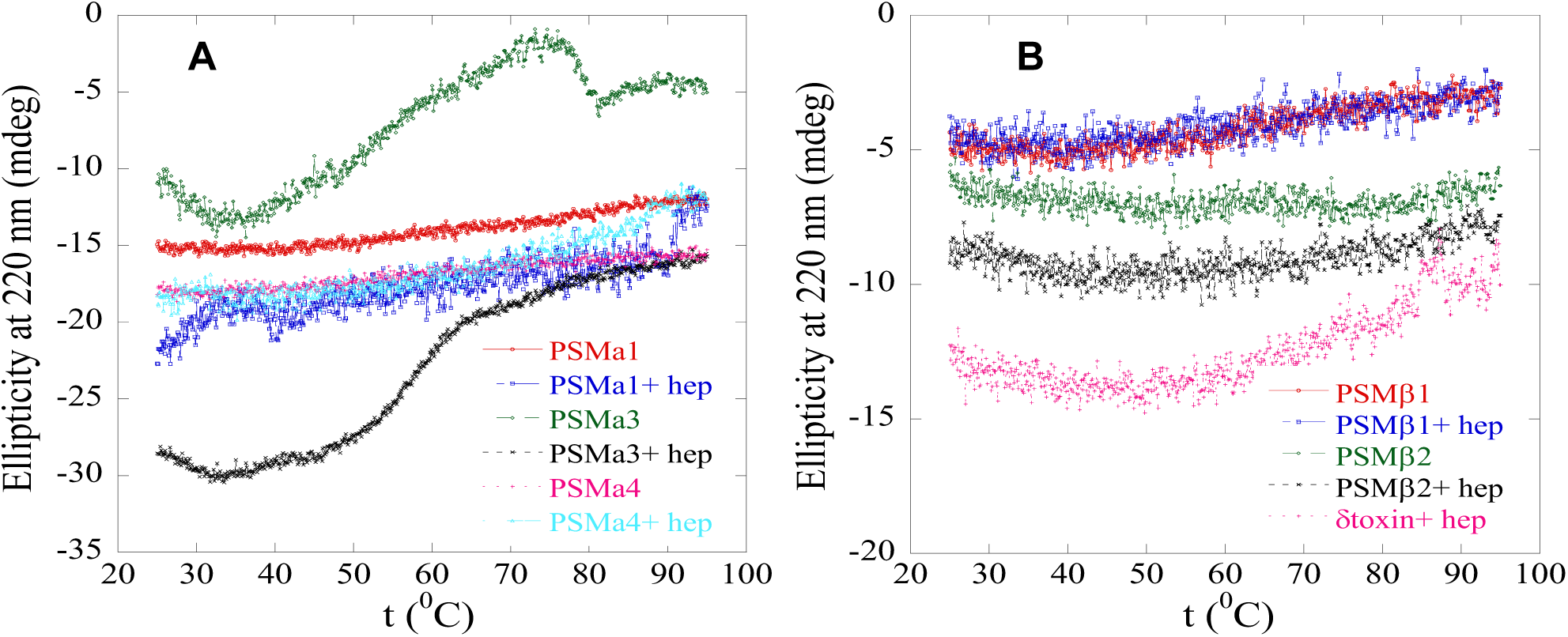
(A) CD thermal scans from 20 to 95°C of α-PSM fibrils incubated in absence and presence of heparin. The concentration of heparin are 3 µg/mL for PSMα1, 40 µg/mL for PSMα3, 50 µg/mL for PSMα4, (B) CD thermal scans from 20 to 95°C of β-PSM and δ-toxin fibrils incubated in absence and presence of heparin (Heparin concentrations: 250 µg/mL for PSMβ1, 1 mg/mL for PSMβ2 and 1 mg/mL for δ-toxin).

**Figure S7:**
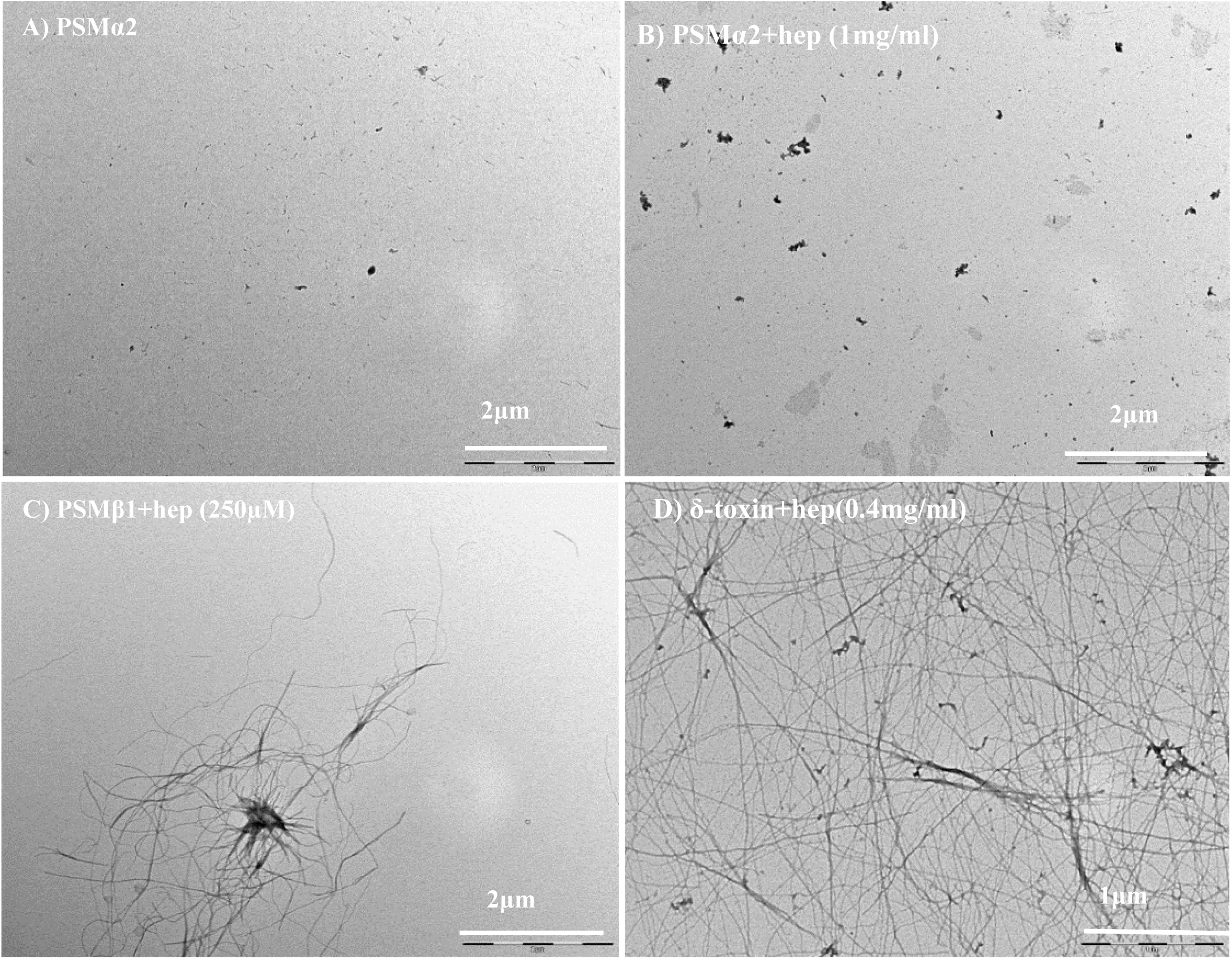
TEM images of PSM fibrils with and without heparin. (a) PSMα2 without heparin, (b) PSMα2 with 1 mg/mL heparin, (c) PSMβ1 with 250 µg/mL heparin and (d) δ-toxin with 0.4 mg/mL heparin. Note that scale bar changes.

**Figure S8.**
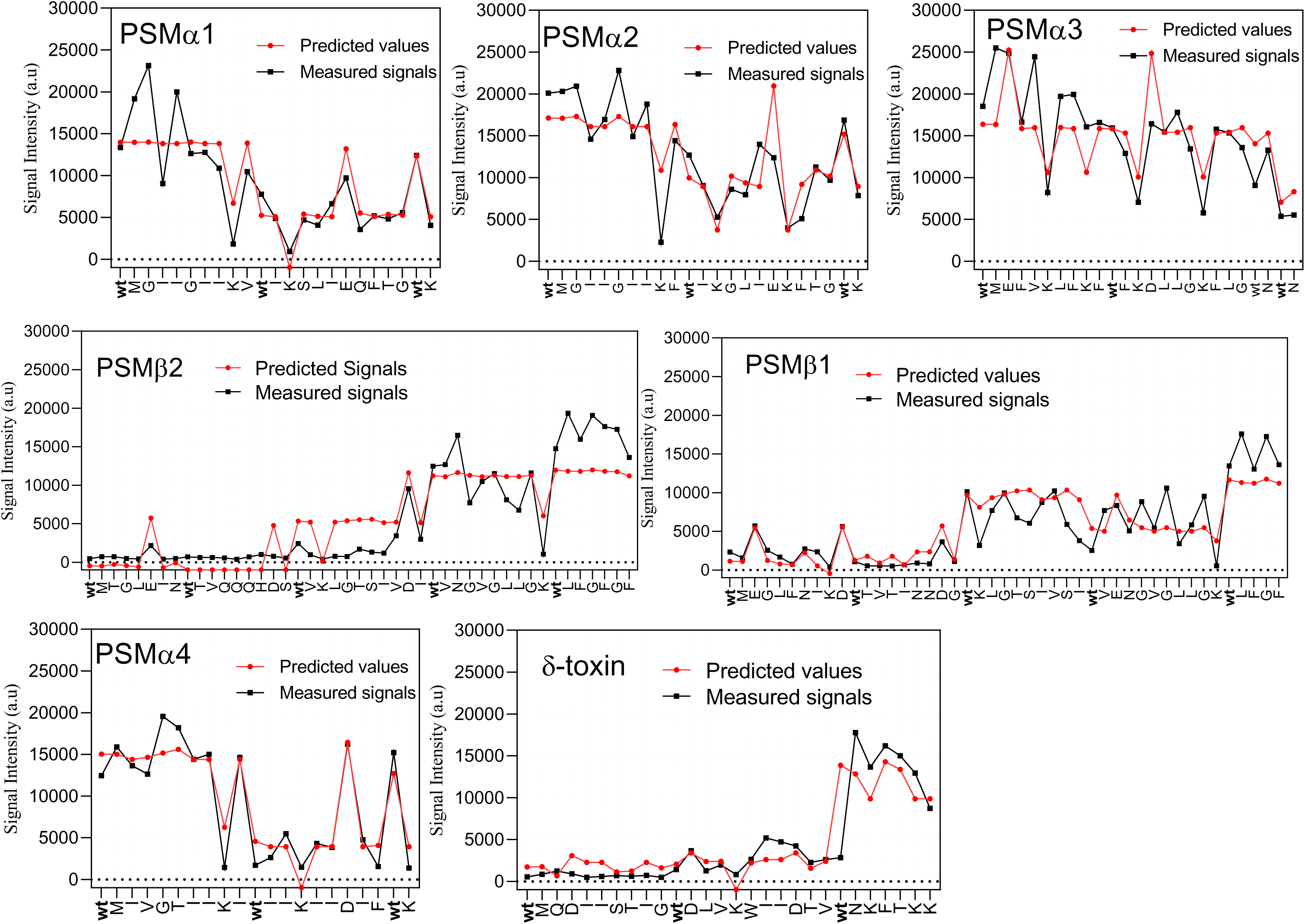
Graphs of predicted and measured signal intensities based on multiple regression with two variables (charge and hydrophobicity) based on data from Ala scan of PSM sequences. Each letter on the x-axis shows the residue that is replaced by Ala.

**Table S1.** Composite rate constant (*k*_+_*k*_n_ in units of conc^-n-1^ time^-2^) of secondary nucleation model from fitting of ThT-data with Amylofit web server.

| Heparin<br>$\mu\text{g/ml}$ | | PSMα1 | Heparin<br>$\mu\text{g/ml}$ | PSMα3 | Heparin<br>$\mu\text{g/ml}$ | PSMβ1 | Heparin<br>$\mu\text{g/ml}$ | PSMβ1 | Heparin<br>$\mu\text{g/m}$ | PSMβ2 |
| --- | --- | --- | --- | --- | --- | --- | --- | --- | --- | --- |
| Individual fits | 0 | $6.84 \times 10^{-10}$ | 0 | 536 | 0 | $3.57 \times 10^{+17}$ | 50 | $1.52 \times 10^{+12}$ | 0 | 362.00 |
| | 0.5 | $1.19 \times 10^{-8}$ | 10 | 900 | 5 | $2.78 \times 10^{+17}$ | 75 | $3.66 \times 10^{+10}$ | 0.1 | 20.00 |
| | 1 | $1.22 \times 10^{-7}$ | 20 | $1.30 \times 10^{+3}$ | 10 | $2.64 \times 10^{+17}$ | 100 | $5.04 \times 10^{+7}$ | 0.15 | 10.60 |
| | 1.5 | $4.43 \times 10^{-6}$ | 25 | $1.75 \times 10^{+3}$ | 20 | $4.66 \times 10^{+17}$ | 150 | $4.09 \times 10^{+3}$ | 0.2 | 9.55 |
| | 2 | $1.63 \times 10^{-4}$ | 30 | $2.64 \times 10^{+3}$ | 30 | $1.19 \times 10^{+18}$ | 200 | $1.57 \times 10^{+5}$ | 0.25 | 6.55 |
| | 3 | $1.32 \times 10^{-3}$ | 40 | $3.75 \times 10^{+3}$ | 40 | $1.87 \times 10^{+18}$ | 250 | $1.32 \times 10^{+4}$ | 0.5 | 0.87 |
|  | - |  | - |  | - |  | - |  | 1 | 0.53 |
| Global fits | $m_0 = 110 \mu\text{M}$<br>$n_c = 1.98 \times 10^{-5}$<br>$k_+k_2 = 2.63 \times 10^{+5}$<br>($\text{conc}^{-n_2-1} \text{ time}^{-2}$)<br>$n_2 = 0.542$<br>MRE:0.00488 | | $m_0 = 96 \mu\text{M}$<br>$n_c = 0.723$<br>$k_+k_2 = 1.57 \times 10^{+8}$<br>($\text{conc}^{-n_2-1} \text{ time}^{-2}$)<br>$n_2 = 0.695$<br>MRE:0.00176 | | $m_0 = 55 \mu\text{M}$<br>$n_c = 4$<br>$k_+k_2 = 2.38 \times 10^{+4}$<br>($\text{conc}^{-n_2-1} \text{ time}^{-2}$)<br>$n_2 = 0.316$<br>MRE:0.00418 | | $m_0 = 55 \mu\text{M}$<br>$n_c = 3.92$<br>$k_+k_2 = 6.71 \times 10^{+5}$<br>($\text{conc}^{-n_2-1} \text{ time}^{-2}$)<br>$n_2 = 0.2$<br>MRE:0.00905 | | $m_0 = 5.61 \mu\text{M}$<br>$n_c = 0.762$<br>$k_+k_2 = 3.05 \times 10^{+4}$<br>($\text{conc}^{-n_2-1} \text{ time}^{-2}$)<br>$n_2 = 0.00378$<br>MRE: 0.0117 | |

